# Bacteria-herpesvirus interaction reveals a proviral role of ISGylation in promoting herpesvirus capsid assembly

**DOI:** 10.64898/2026.01.19.700396

**Authors:** Shutong Li, Shu Feng, Yongzhen Liu, Chao Qin, Jianglin Zhang, Takeru Saito, Xinchi Xie, Wen Fu, Rui Wang, Yue Zheng, Zhenfeng Shu, Wayne Yeh, Sean Sasaki, Natalia Tjokro, Lin Chen, Joshua Munger, Jae Jung, Casey Chen, Pinghui Feng

## Abstract

Herpesviruses are common pathogens of the oral cavity, yet how they interact with other oral microbes are poorly understood. Using murine gamma-herpesvirus 68 (MHV68) as a model for human gamma-herpesviruses, we discover that ISGylation is hijacked to facilitate viral capsid assembly and lytic replication. Co-infection with the oral *Aggregatibacter actinomycetemcomitans* (*A.a*.) and MHV68 synergistically induced interferon-stimulated gene 15 (ISG15) expression and global ISGylation. Proteomic profiling revealed viral structural proteins as the dominant targets of ISGylation in MHV68-infected cells. Genetic ablation of ISGylation, using ISGylation-resistant recombinant MHV68 and ISG15-deficient mouse embryonic fibroblasts (MEFs), demonstrated that ISGylation of the major capsid protein ORF25 is required for efficient capsid assembly and maturation into the infectious C-type virions. Strikingly, introduction of a *de novo* ISGylation site into the ISGylation-resistant MHV68 was sufficient to restore ISGylation, capsid assembly, and virion maturation. Consistent with these findings, *A.a.* failed to enhance MHV68 lytic replication in ISG15-deficient MEFs. Extending this mechanism to representative herpesviruses, we showed that ISGylation of major capsid proteins is broadly required for efficient lytic replication. Together, these findings uncover a previously unrecognized strategy by which herpesviruses exploit an ISG15 innate immune effector, amplified by microbial co-infection, to promote virion biogenesis and productive infection.

## Introduction

Herpesviruses are ubiquitous in humans and cause significant morbidity and mortality in immune-compromised individuals, such as patients with acquired immunodeficiency syndrome (AIDS)^2^. Human herpesvirus 8 (HHV-8), also known as Kaposi’s sarcoma-associated herpesvirus (KSHV), is the etiological agent responsible for Kaposi’s sarcoma (KS) and two rare forms of lymphoma, i.e., primary effusion lymphoma (PEL) and multicentric Castleman’s disease (MCD)^3, 4, 5^. The oral cavity provides an environment conducive for the infection, replication, transmission, and pathogenesis of herpesviruses^1,2,6^. Piling evidence suggests that bacterial infection is associated with the increased KSHV load in the oral cavity^20, 21^. We previously reported that ongoing oral bacterial infection can stimulate KSHV lytic replication and identified specific bacterial species, particularly *Neisseria gonorrhoeae* and *Aggregatibacter actinomycetemcomitans (A.a.)*, that stimulate KSHV replication^22^. During the late stages of lytic replication, herpesviruses express a large array of proteins that are packaged into the progeny virion in several artificially defined processes, including capsid assembly, tegumentation and maturation/egress^7^. However, our understanding in the regulatory mechanisms governing these herpesviral replication processes remains rudimentary at best, partly due to the limited availability of tractable experimental systems.

Interferon-stimulated gene 15 (ISG15) is a ubiquitin-like pleiotropic protein and one of the most highly expressed ISGs^10^. Intracellularly, ISG15 regulates protein translation, stability, trafficking, cytokine production, and immune response by covalently modifying host or viral proteins through a post-translational process termed ISGylation^11, 12^. For example, ISGylation of the influenza A virus non-structural protein 1 (NS1) disrupts its dimerization and nuclear translocation, thereby attenuating the NS1-mediated interferon (IFN) antagonism^13, 14^. Similarly, ISGylation of the nucleoprotein (NP) of influenza B virus interferes with the NP oligomerization and viral ribonucleoprotein (vRNP) complex assembly, ultimately impairing viral replication^15^. Beyond direct antiviral effects, ISGylation also modulates host immune response. ISGylation has been shown to negatively regulate the retinoic acid-inducible gene I (RIG-I)-mediated antiviral signaling and promote viral replication^16^. Moreover, ISGylation can stabilize the de-ISGylating enzyme USP18 by preventing its proteasomal degradation, which dampens the JAK-STAT signaling and interferon responses^17^. The functional consequences of ISGylation are highly context- and substrate-dependent and can either restrict or facilitate microbial infection. Though the potential proviral roles of ISGylation are emerging, the molecular underpinnings remain poorly defined.

Murine gammaherpesvirus 68 (MHV68), which is genetically related to human gammaherpesviruses including KSHV and Epstein-Barr virus (EBV), infects a wide range of mouse strains and provides a well-established model for studying the molecular event relevant to human gammaherpesvirus infection^18,19^. The robust lytic replication of MHV68 after *de novo* infection enables the biochemical characterization of the late-stage processes of replication possible, which is very challenging for human KSHV and EBV. Here, we show that *A.a.* promotes MHV68 lytic replication. Interestingly, *A.a.* and MHV68 synergistically induce ISGylation, and ISGylation of the major capsid protein ORF25 promotes the capsid and virion assembly of MHV68, which can be extended to other representative herpesviruses. Findings from this study uncover an unexpected proviral role of ISGylation in promoting virion assembly during herpesvirus lytic replication, providing a novel system to dissect the molecular events in bacteria-virus infection.

## Results

### Oral bacteria stimulate MHV68 lytic replication

We have previously reported that oral bacteria enhance the lytic replication of human KSHV^22^, the etiological agent of KS, PEL and MCD^23^. However, KSHV has limited replication after *de novo* infection, making it difficult to biochemically characterize the molecular events of its lytic replication. The closely related MHV68 undergoes robust lytic replication in mouse fibroblasts (e.g., mouse embryonic fibroblasts [MEFs]), providing a useful system to model human KSHV and EBV. Thus, we tested whether oral *A.a.* can promote MHV68 lytic replication. To effectively monitor MHV68 infection, we used a recombinant MHV68 carrying a GFP at the K3 locus, designated as MHV68 K3/GFP (Fig. 1a). When MEFs were infected with MHV68 followed by *A.a.* infection, we found that *A.a.* infection elevated the replication of MHV68 K3/GFP as indicated by more robust GFP fluorescence than MHV68 K3/GFP without *A.a.* (Fig. 1a). Consistent with this result, plaque assay using medium containing extracellular virions and infected cell lysates showed that *A.a.* infection elevated the infectious MHV68 K3/GFP by ∼4-6-fold (Fig. 1b and Extended Data Fig. 1a). These results show that *A.a.* infection promotes MHV68 lytic replication.

**Figure 1.**
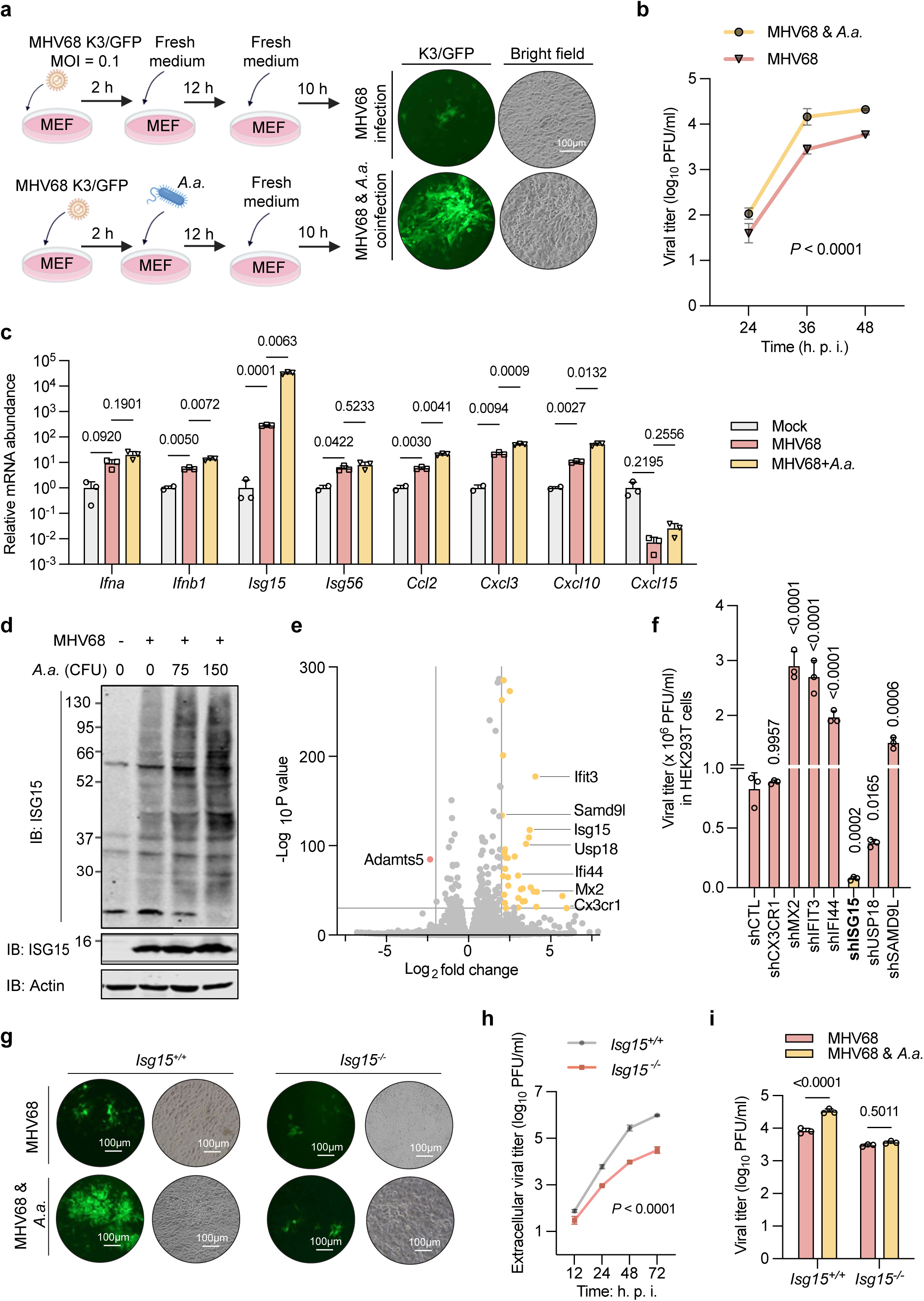
*Aggregatibacter actinomycetemcomitans (A.a.)* promotes MHV68 lytic replication. **a-c**, Schematic illustration of the procedure to coinfect mouse embryonic fibroblasts (MEFs) with MHV68 K3/GFP and *A.a.* (D7SS). MEFs were infected with MHV68 K3/GFP (MOI = 0.1, 2 hpi), with or without *A.a.* (75 CFU/cell, 12 hpi), washed with DMEM to remove *A.a.,* cultured with fresh medium, and recorded with a fluorescence microscope at 24 h after MHV68 infection. Representative images were shown. Scale bars, 100 μm (**a**). MHV68 titer in the medium at indicated time points was determined by plaque assay using NIH 3T12 monolayer (**b**). Total RNA was extracted, reverse-transcribed, and analyzed by real-time PCR with primers specific for *Ifna*, *Ifnb1*, *Isg15*, *Isg56*, *Ccl2*, *Cxcl3*, *Cxcl10*, and *Cxcl15* (**c**). **d**, Whole cell lysates (WCLs) of mock-, MHV68-, and MHV68 and *A.a.*-infected MEFs were analyzed by SDS-PAGE and immunoblotting with indicated antibodies. **e**, Volcano plot showing differential gene expression in mock- and MHV68-infected NIH 3T3 cells as determined by RNA sequencing. The significance and fold-change were converted to −Log_10_ (*P*-value) and Log_2_ (fold-change), respectively. The vertical and horizontal lines show the cut-off of fold-change = ± 8, and of *P*-value = 0.0005, respectively. Yellow dots represent up-regulated genes (> 8-fold, *P* < 0.0005). Seven interferon-stimulated genes (ISGs) (*Ifit3*, *Samd9l*, *Isg15*, *Usp18*, *Ifi44*, *Mx2*, and *Cx3cr1*) with significant up-regulation were labelled. **f**, The effect of individual ISG on MHV68 lytic replication was determined by plaque assay using ISG-depleted HEK293T cells infected with MHV68 (MOI = 0.5). **g** and **h**, *Isg15^+/+^* and *Isg15^-/-^* MEFs were infected with MHV68 (MOI = 0.5), with or without *A.a.,* and analyzed with a fluorescence microscope. Representative images were shown (**g**). Virus-containing medium was analyzed by plaque assay (**h**). **i**, Multi-step growth curve of MHV68 (MOI = 0.01) in the medium of *Isg15^+/+^*and *Isg15^-/-^*MEFs was characterized by plaque assay. Data represent three independent experiments.

### ISG15 expression promotes MHV68 lytic replication

To assess whether innate immune response is altered by *A.a.* co-infection, we profiled the expression of representative inflammatory genes, including IFNs, ISGs, and chemokines. In general, MHV68 K3/GFP infection modestly induced the expression of these inflammatory genes, which was further enhanced by *A.a.* co-infection (Fig. 1c). Notably, the expression of *Isg15* was induced by MHV68 K3/GFP by >100-fold, and *A.a.* coinfection further elevated by another ∼100-fold. By contrast, the expression of *Cxcl15* was reduced by MHV68 K3/GFP, with and without *A.a.* infection. We have an outstanding interest in understanding the regulatory roles of protein post-translational modifications, we thus further examined whether ISG15 expression is important for MHV68 infection. Indeed, immunoblotting analysis showed that MHV68 infection greatly elevated ISG15 expression and the smeared signal of ISG15, with the latter being ISGylated proteins including poly-ISG15 chains (Fig. 1d). The induction of ISG15 and ISGylation was further enhanced by *A.a.* infection. Thus, MHV68 infection induces ISG15 expression and likely protein ISGylation.

To determine the role of ISG15 expression in MHV68 infection, we annotated a previous published gene expression of MHV68-infected cells, focusing on the ISGs^24^. This identified seven ISGs whose expression was induced by > 8-fold by MHV68 infection (Fig. 1e). We then depleted these genes in HEK293T cells with shRNA targeting these genes (Extended Data Fig. 1b) and examined their effect on MHV68 lytic replication. As expected, the knockdown of four out of seven ISGs modestly increased MHV68 lytic replication, while knockdown of CXCR1 had marginal effect on MHV68 lytic replication (Fig. 1f). Surprisingly, the knockdown of ISG15 and, to a less extent, USP18 greatly reduced MHV68 lytic replication. Next, we compared the replication kinetics of MHV68 in wildtype and *Isg15^-/-^* MEFs. We found that ISG15 deficiency reduced MHV68 lytic replication as indicated by GFP fluorescence and plaque assay determining the extracellular infectious virions and cell-associated virions (Fig. 1g, 1h and Extended Data Fig. 1c). Furthermore, *A.a.* co-infection elevated MHV68 extracellular virions in wildtype MEFs, but failed to do so in *Isg15^-/-^* MEFs (Fig. 1i). Taken together, these results collectively support the conclusion that ISG15 expression promotes MHV68 lytic replication and *A.a.* stimulates MHV68 lytic replication in an ISG15-dependent manner.

### ISG15 promotes the virion assemble of MHV68 lytic replication

To probe the role of ISG15 in MHV68 lytic replication, we assessed the key steps of lytic replication, including the entry, trafficking to the nucleus, mRNA expression, genome replication, and virion released into the extracellular compartment using wildtype and *Isg15^-/-^* MEFs for MHV68 infection at MOI of 5 (Fig. 2a). For MHV68 entry, we examined the intracellular viral genome by real-time PCR at 30 and 90 min post-infection, which showed that loss of ISG15 had no effect on viral entry (Fig. 2b). To analyze the trafficking of MHV68 nucleocapsid to the nucleus, we isolated the cytosolic and nuclear fractions via subcellular fractionation. At 1 and 2 h post-infection, the nuclear MHV68 genome were identical in wildtype and *Isg15^-/-^* MEFs as analyzed by real-time PCR (Fig. 2c). There were slightly lower levels of the MHV68 genome in the nuclear and cytoplasm of *Isg15^-/-^* MEFs than those in wildtype MEFs, which is likely due to new genome replication. Interestingly, when viral RNA was analyzed by reverse transcription and real-time PCR, the mRNA levels of MHV68 lytic genes were slightly higher in *Isg15^-/-^* MEFs than wildtype MEFs that were infected with MHV68 (Fig. 2d, Extended Data Fig. 2a-f). At 12 hpi, the copy number of the MHV68 genome was slightly lower (<1.5-fold) in *Isg15^-/-^*MEFs than wildtype MEFs (Fig. 2e). These results show that loss of ISG15 expression slightly but differentially altered viral gene expression and genome replication of MHV68 in MEFs.

**Figure 2.**
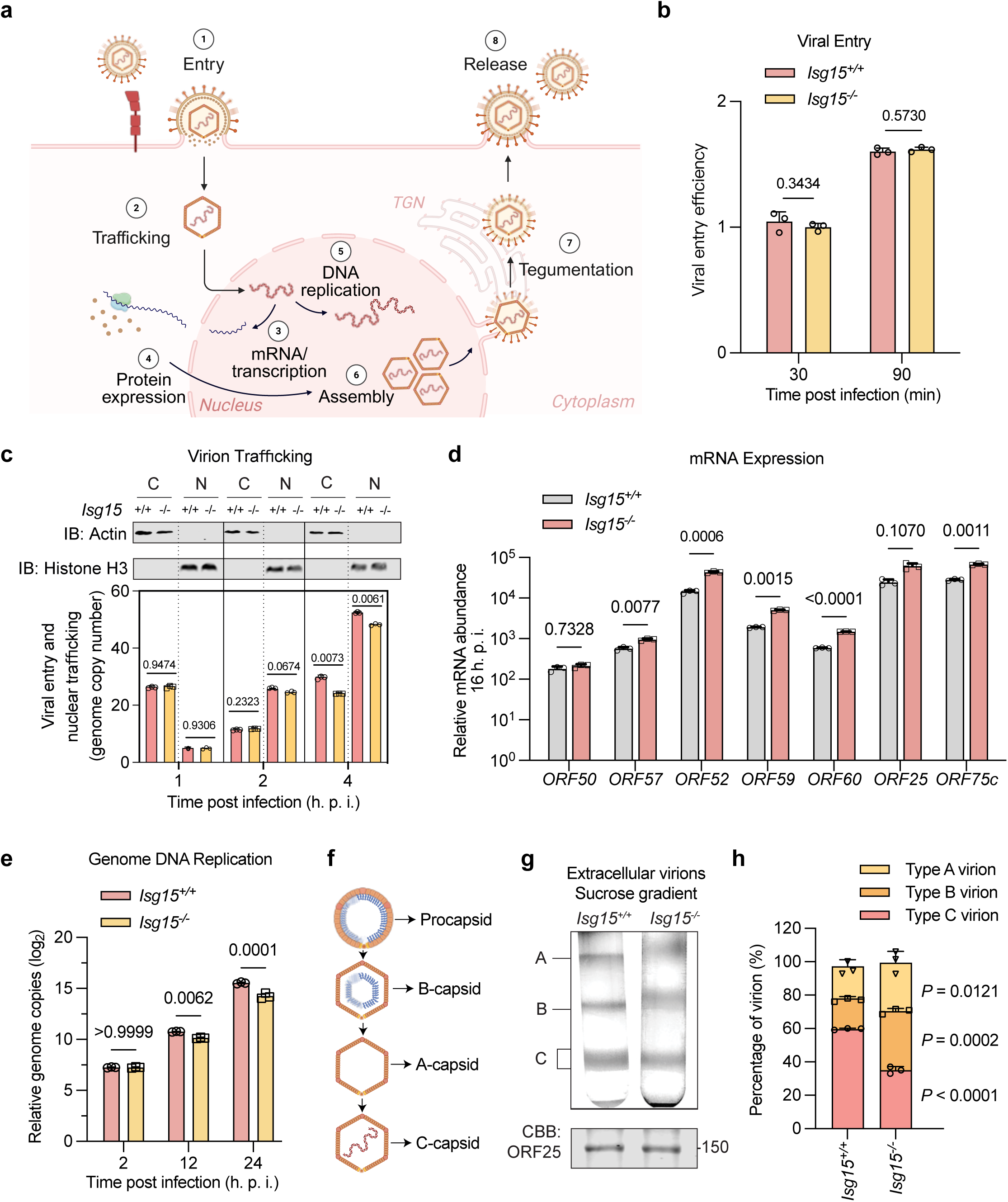
ISG15 is required for the virion assembly of MHV68 lytic replication. **a**, Schematic illustration of key steps of MHV68 lytic replication: 1) viral entry (attachment and membrane fusion); 2) capsid trafficking to the nucleus; 3) viral transcription; 4) protein synthesis; 5) viral DNA genome replication; 6) capsid and virion assembly; 7) tegumentation; and 8) maturation and egress. Capsid assembly, tegumentation, maturation and egress are the late-stage steps. **b**, Entry analysis of MHV68 in *Isg15^+/+^* and *Isg15^-/-^* MEFs. Cells were infected with MHV68 (MOI = 5), and the intracellular genome in total DNA at the indicated time point was analyzed by real-time PCR. **c**, Trafficking analysis of MHV68 in *Isg15^+/+^* and *Isg15^-/-^* MEFs. MHV68-infected cells were subjected to sequential centrifugation to obtain the cytosolic (C) and nuclear (N) fraction. The genome from each fraction was analyzed by real-time PCR. Cytosolic and nuclear fraction were analyzed by SDS-PAGE and immunoblotting with indicated antibodies. **d**, Transcription of the MHV68 genome in *Isg15^+/+^* and *Isg15^-/-^* MEFs at 16 hpi was analyzed by real-time PCR with gene-specific primers after reverse transcription. Two immediate-early, early, and late genes were examined the MHV68 transcription program. **e**, MHV68 genome replication analysis. Total intracellular DNA was extracted from MHV68-infected *Isg15^+/+^* and *Isg15^-/-^* MEFs and subjected to real-time PCR to measure viral genome copy number at indicated time points. **f**, Schematic illustration of capsid assembly pathway. The assembly of herpesvirus capsid starts with the formation of a procapsid, which is supported by scaffolding proteins. As maturation proceeds, scaffolding proteins are cleaved and degraded, and the capsid transitions from a round shape to an icosahedral structure. Concomitantly, the viral DNA is packaged into the capsid, forming the C-capsid. Failure in DNA packaging produces A-capsids that are empty, or B-capsids that still contain remnants of scaffolding proteins. Blue: scaffolding protein; Orange: major capsid protein. **g,h**, Virion assembly analysis of MHV68 in *Isg15^+/+^*and *Isg15^-/-^*MEFs. Extracellular MHV68 virions in the medium were collected, concentrated by ultracentrifugation, and resolved by sucrose gradient centrifugation. The virion bands were photographed (**g**) and quantified by ImageJ (**h**). Data represent three-independent experiments.

Next, we assessed the late stages of MHV68 lytic replication, specifically the three types of extracellular virions according to their capsids produced from wildtype and *Isg15*^-/-^ MEFs (Fig. 2f and Extended Data Fig. 2g). While B-type virions contain scaffold proteins within the capsid, A-type virions have empty capsids. The C-type virions are packaged with the viral genome and thus infectious^7^. When extracellular MHV68 virions were analyzed by sucrose gradient centrifugation, we found that virions produced from *Isg15^-/-^*MEFs had significantly higher percentage of A- and B-type virions, which was supported by semiquantitative measurement of the band density (Fig. 2g and h). Correspondingly, the C-type virions were significantly lower from *Isg15^-/-^*MEFs than those from wildtype MEFs. These results show that loss of ISG15 impedes the virion assembly likely at the capsid assembly and genome package.

### Identification of ISGylated proteins of MHV68

Considering the proviral roles of ISG15 and the smearing phenotype of ISG15 signal in MHV68-infected cells, we hypothesize that ISG15 is conjugated to viral, and likely cellular, proteins to promote MHV68 lytic replication. To identify ISGylated proteins, we established *Isg15^-/-^* MEFs that stably express FLAG-ISG15 via lentiviral infection for MHV68 infection (Extended Data Fig. 1a-c). To avoid non-covalent binding of ISG15, we denatured the lysates of MHV68-infected MEFs via boiling at ∼100 °C for 10 minutes in lysis buffer containing 1% SDS and diluted it 10-fold for immunoprecipitation with anti-FLAG M2 agarose (Fig. 3a). Precipitated proteins were subjected to tandem mass spectrometry to identify peptides that carry a diglycine (GG) motif, the remnant of trypsin digestion of conjugated ubiquitin, ISG15 or SUMO (Fig. 3b). This analysis identified a number of peptides carrying the GG motif, which were matched to MHV68 proteins (Extended Data Fig. 3d). The cellular proteins with peptide carrying the GG motif were identified and will be reported in the future. Surprisingly, most of these MHV68 proteins are structural proteins, including major capsid protein ORF25, small capsid protein ORF65, triplex capsid protein 2, portal protein ORF43, capsid assembly protein ORF62, tegument protein ORF52, ORF75b, ORF75c, and DNA replication processivity factor ORF59 (Extended Data Fig. 3e). This result suggests that virion structural proteins are the major targets of ISGylation in MHV68 infection. To determine whether these virion structural proteins undergo ISGylation, we analyzed precipitated MHV68 proteins with antibody to ISG15 with denatured whole cell lysates of transfected HEK293T cells. To facilitate ISGylation, E1 (UBE1L) and E2 (UBE2) enzymes were over-expressed in the same transfection. As shown in Extended Data Fig. 3f, the ISG15 signal was readily detected with precipitated MHV68 proteins, including ORF25, ORF43, ORF65 and ORF75c. Given the highest ratio of GG-modified peptides (Fig. 3c) and the crucial role of ORF25 major capsid protein in virion assembly and lytic replication, we further characterized the ISGylation of ORF25 during MHV68 infection. To further confirm the ISGylation of ORF25, we infected MEFs with a recombinant MHV68 carrying the FLAG epitope at the C-terminus of ORF25 and precipitated ORF25 for immunoblotting analysis. This clearly identified a single distinct species of ORF25 that approximately corresponds to the ORF25 with two units of ISG15 based on its apparent molecular weight (Fig. 3d). These results conclude that virion structural proteins are major targets of ISGylation during MHV68 infection.

**Figure 3.**
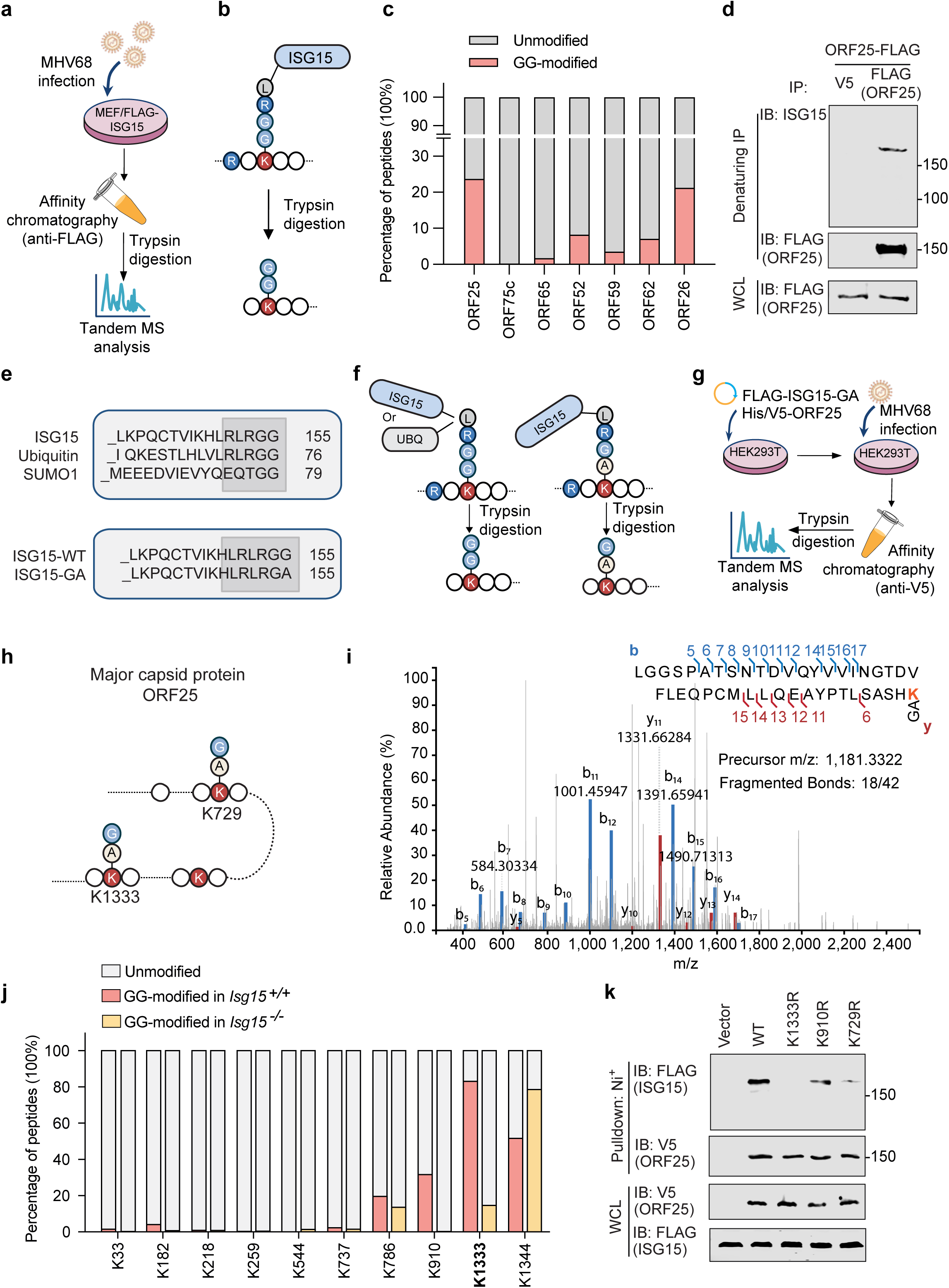
Identification of MHV68 proteins with ISGylation. **a-c**, ISG15-conjugated-proteomic analysis: *Isg15^-/-^* MEFs reconstituted with FLAG-ISG15 were infected with MHV68 (MOI=2, 48 hpi). Affinity purification of ISG15 under denaturing condition was performed followed by tandem mass spectrometry analyses to identify proteins/peptides that carry a diglycine (GG) motif (**a**). Schematic illustration of the GG-containing peptide derived from ISGylated proteins. (**b**). Quantification of MHV68 proteins by the abundance of GG-modified peptides and the total peptides (**c**). **d**, MEFs was infected with recombinant MHV68.ORF25-FLAG (MOI=2, 48 hpi). ORF25 was purified with anti-FLAG M2 agarose under denaturing condition and subjected to immunoblotting with indicated antibodies. **e**, Top: amino acid sequence alignment of ISG15, ubiquitin, and SUMO1. Bottom: amino acid sequence alignment of ISG15-WT and ISG15-G155A. **f**, Schematic illustration of GA-modified peptides generated by trypsin digestion of ISGylated proteins in cells expressing ISG15-G155A. **g** and **h**, Identification of ORF25 ISGylation site using the ISG15-G155A mutant: HEK293T cells were co-transfected with FLAG-ISG15-G155A and ORF25-His/V5, followed by MHV68 infection. Affinity purification of ORF25 under denaturing condition followed by tandem mass spectrometry analyses identified peptides containing K729 and K1333 (**h**). **i**, The m/z spectrum of the peptide containing K1333 that carries the GA remnant is shown and is highlighted in red. **j**, ORF25 was purified from *Isg15^+/+^* and *Isg15^-/-^* MEFs infected with MHV68.ORF25-FLAG (MOI=2, 48 hpi) and analyzed by LC-MS/MS to quantitatively determine ORF25 ISGylation sites. ISGylation efficiency is the ratio of GG-modified peptide abundance to the total peptide abundance specific for a site. **k**, HEK293T cells were co-transfected with FLAG-ISG15, ORF25-His/V5 or the mutants (K1333R, K729R, and K910R), E1, and E2, followed by immunoprecipitation under denaturing condition and immunoblotting with indicated antibodies.

Cellular ISG15, ubiquitin and SUMO share the same carboxyl-terminus that leave a GG motif on peptides after trypsin digestion (Fig. 3e). To further differentiate the GG motif is derived from ISGylation rather than ubiquitination or SUMOylation, we generated an ISG15-G155A mutant that will leave a GA on trypsinized peptides (Fig. 3f). In transfected HEK293T cells, the GA mutant was efficiently conjugated to ORF25 compared to wildtype ISG15 (Extended Data Fig. 3g). We then established HEK293T cells stably expressing FLAG-ISG15-GA mutant. These cells were transfected with a plasmid expressing 6×His- and V5-tagged ORF25 and infected with MHV68 to promote ISGylation. Affinity purified ORF25 was subjected to tandem MS analysis to identify GA-containing peptides (Fig. 3g). This confirmed K1333 and K729 as the ISGylation sites (Fig. 3h, i, and Extended Data Fig. 3h). To further characterize the ISGylation sites, we compared the site modification of ORF25 by GG in wildtype and *Isg15^-/-^* MEFs infected with recombinant MHV68 expressing FLAG-ORF25. Tandem MS analysis using purified ORF25 showed that >80% of peptides containing K1333 were modified by diglycine in wildtype MEFs, and additional sites include K1344, K910 and K786, were modified to various degrees (Fig. 3j and Extended Data Fig. 3i). Remarkably, loss of ISG15 expression reduced the percentage of the peptide containing diglycine at K1333 to <20% and that of the peptide containing diglycine at K910 from ∼35% to base line (Fig. 3j). Interestingly, ISG15 deficiency elevated the percentage of peptide containing diglycine at K1344, indicating a possible compensatory mechanism via ubiquitination or SUMOylation. These results nominate K1333, K910 and K729 as major ISGylation sites of ORF25. To delineate the contribution of individual residues, we generated ORF25 constructs containing individual K to R mutations at K1333, K910 and K729 for ISGylation analysis. Immunoprecipitation under denaturing condition followed by immunoblotting analysis showed that the K1333R abolished, while K729R and, to a less extent, K910, greatly reduced ORF25 ISGylation in transfected HEK293T cells (Fig. 3k). These results conclude that ORF25 is ISGylated at multiple lysine residues.

### Defects in virion assembly of recombinant MHV68 deficient in ORF25 ISGylation

To characterize the role of ORF25 ISGylation, we generated recombinant MHV68 using the artificial bacterial chromosome (BAC) system^25^. Considering that the K1333R mutation completely abolished ORF25 ISGylation, we engineered recombinant MHV68 containing ORF25-K1333R (Extended Data Fig. 4a). Multi-step growth curves of recombinant MHV68 showed that the K1333R mutation reduced both the extracellular and intracellular titer of MHV68 (Fig. 4a and Extended Data Fig. 4b), while K910R and K729R mutations marginally reduced viral titer (Extended Data Fig. 4c). The lower intracellular titer of MHV68.ORF25-K1333R is likely due to the amplification of the multi-step growth process of MHV68 infection. Additionally, MHV68 ORF25-K1333R also formed smaller plaques on NIH 3T12 monolayer (Fig. 4b). With recombinant MHV68 carrying ORF25-FLAG and ORF25-K1333R-FLAG, we demonstrated that the K1333R mutation abolished ORF25 ISGylation in MEFs infected MHV68 (Fig. 4c), which agrees with the result from transfected HEK293T cells (Fig. 3k). When tested with co-infection of *A.a*., we found that *A.a.* co-infection failed to enhance MHV68.ORF25-K1333R, but did so for wildtype MHV68 (Fig. 4d). These results show that ORF25 ISGylation is important for MHV68 lytic replication.

**Figure 4.**
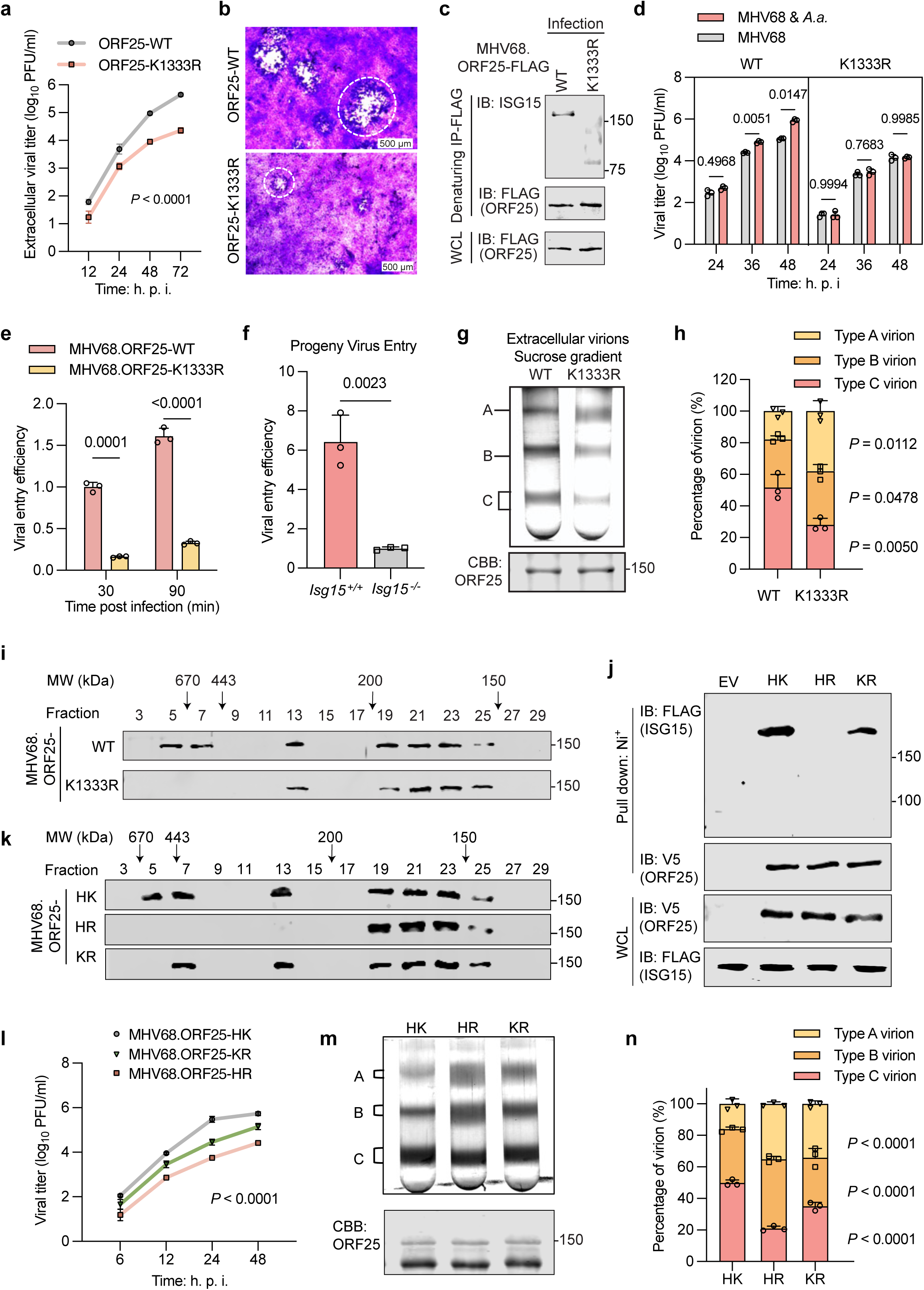
ISGylation of ORF25 at K1333 is required for efficient MHV68 lytic replication. Recombinant MHV68 containing ISGylation-resistant K1333R mutation in ORF25 (MHV68.ORF25-K1333R) was generated by BAC. **a,b,** Multi-step growth curve of MHV68.ORF25-WT and MHV68.ORF25-K1333R (MOI=0.01) was characterized by plaque assay using extracellular virions in the medium (**a**). Viral input was normalized by packaged genome copy number. Representative plaques were shown (**b**). **c**, MEFs were infected with indicated recombinant MHV68 (MOI=2, 48 hpi). ORF25 was purified with anti-FLAG under denaturing condition and subjected to immunoblotting with indicated antibodies. **d,** MEFs were infected with MHV68.ORF25-WT and MHV68.ORF25-K1333R (MOI=0.1) for 2 h, coinfection and plaque assay were performed to determine extracellular viral titer as described in Fig. 1a. **e**, Entry analysis of MHV68.ORF25-WT and MHV68.ORF25-K1333R. MEFs were infected with indicated recombinant MHV68 (MOI = 5), and the intracellular genome at the indicated time point was analyzed by real-time PCR using total DNA. **f**, Entry analysis of the progeny virus from *Isg15^+/+^* and *Isg15*^-/-^ MEFs. *Isg15^+/+^*and *Isg15^-/-^*MEFs were infected with MHV68 (MOI = 0.1 for 72 hpi). Progeny virus was normalized with the packaged genome copy number and used to infect target cells (MOI = 5). **g**,**h**, Virion assembly analysis of MHV68.ORF25-WT and MHV68.ORF25-K1333R. MEFs were infected with indicated recombinant MHV68 normalized with equal entry. Extracellular virions in the medium were collected, concentrated by ultracentrifugation, and resolved by sucrose gradient centrifugation. The virion bands were photographed (**g**) and quantified by ImageJ (**h**). **i**, ORF25 proteins were purified from MEFs infected with indicated recombinant MHV68 (MOI=2, 48 hpi) with anti-FLAG M2 agarose and analyzed by size exclusion chromatography. Fractions (0.8 mL) were concentrated and analyzed by immunoblotting with anti-FLAG. Numbers on the top indicate the size of molecular weight. **j,k**, Recombinant MHV68 containing H1332K mutation in ORF25-K1333R (MHV68.ORF25-KR) was generated by BAC. MEFs was infected with MHV68.ORF25-WT (WT), MHV68.ORF25-K1333R (HR), and MHV68.ORF25-H1332K/K1333R (KR) (MOI=2, 48 hpi). ORF25 was purified with anti-FLAG M2 agarose under denaturing condition and subjected to immunoblotting with indicated antibodies (**j**). Alternatively, purified ORF25 was eluted and subjected to size exclusion chromatography (**k**). **I,** Multi-step growth curve of indicated recombinant virus (MOI=0.01) in MEFs was characterized by plaque assay using extracellular virions in the medium. Viral input was normalized by the packaged genome copy number. **m**,**n**, Virion assembly analysis of indicated recombinant virus was performed. The virion bands were photographed (**m**) and quantified by ImageJ (**n**).

To further define the steps of MHV68 lytic replication that ORF25 ISGylation is required, we analyzed the key steps of lytic replication as shown in Fig. 2a. Entry analysis showed that MHV68.ORF25-K1333R entered cells at ∼15% efficiency of wildtype MHV68 when intracellular viral genome was quantified by real-time PCR (Fig. 4e). Importantly, loss of ISG15 also reduced the entry efficiency of the extracellular virions produced from MEFs (Fig. 4f and Extended Data Fig. 4d), consistent with the notion that ISGylation of virion structural proteins contributes to the infectivity of virion progeny. By contrast, viral lytic gene expression was only marginally altered by the ORF25 K1333R mutation (Extended Data Fig. 4e). The ORF25 K1333R slightly reduced the genome replication of MHV68 (Extended Data Fig. 4f). Consistent with this, the ORF25 K1333R mutation reduced the extracellular and intracellular titer of MHV68 produced from MEFs, with more significant effect on extracellular MHV68 titer (Extended Data Fig. 4g). When extracellular MHV68 virions were analyzed by sucrose gradient centrifugation, we found that recombinant MHV68.ORF25-K1333R virus accumulated significantly more A- and B-type virions, with correspondingly lower percentage of C-type virions (Fig. 4g and h). This result recapitulates the phenotype of ISG15-deficiency on extracellular MHV68 virions.

### ISGylation promotes the oligomerization of major capsid protein and capsid assembly

Capsid assembly is one of the main checkpoints in herpesvirus productive infection. We therefore tested whether ISGylation is necessary for the oligomerization of major capsid protein and capsid assembly. When ORF25 was purified from MEFs infected with recombinant MHV68.ORF25-FLAG and analyzed by size exclusion chromatography, we found that wildtype ORF25 eluted as three main species that correspond to the monomer, dimer, and pentamer/hexamer (Fig. 4i). In stark contrast, the ORF25-K1333R mutant eluted primarily as monomer and a minor portion of dimer, with no higher molecular oligomer detected under the same conditions (Fig. 4i). This result show that the mutation abolishing ORF25 ISGylation nullifies the ability of ORF25 to form pentamer or hexamer. Within the MHV68 ORF25, we noted that a histidine residue proceeds the ISGylated lysine, K1333 (Extended Data Fig. 4h). We were curious whether a histidine to lysine mutation will be, at least partly, able to restore ISGylation and capsid assembly given the close proximity of the K1332 and K1333. We then generated a new mutant containing H1332K and K1333R, designated KR mutant (Extended Data Fig. 4h and i). We named the wildtype HK, and the K1333R mutant HR, which were included for subsequent analyses. We first assessed whether the KR mutant was ISGylated compared to the HK wildtype and loss-of-function HR mutant. Precipitation under denaturing condition followed by immunoblotting showed that the KR mutant was ISGylated compared to the HR mutant, but to a less extent than the wildtype HK (Fig. 4j). Consistent with this result, the ORF25-KR mutant restored the formation of dimers and the smaller size of the pentamer/hexamer population (Fig. 4k). These results show that ISGylation is necessary and sufficient for ORF25 oligomerization.

Next, we tested whether the H1332K mutation can restore virion assembly and lytic replication of MHV68. When viral lytic replication was examined by multi-step growth curve using extracellular infectious virions, we found that the H1332K mutantion improved MHV68 lytic replication by >50% (Fig. 4l). Moreover, the KR mutant also increased the C-type virions and correspondingly reduced the A- and B-type virions (Fig. 4m and n). The reduction in B-type virion was much more pronounced than that in A-type virions, indicating the specific and partial effect of the H1332K mutation. Taken together, these results unambiguously demonstrate that ISGylation of ORF25 can promote its oligomerization and capsid formation during MHV68 lytic replication.

### Mouse HERC6 and human HERC5 serve as the E3 ligase of the MHV68 ORF25

It’s previously reported that human HERC5 or its mouse HERC6 homologue acts as an E3 ligase to conjugate ISG15 to target proteins^26^. We first tested whether ORF25 interacts with human HERC5. Indeed, ORF25 readily co-precipitated with endogenous HERC6 in MEFs infected with MHV68.ORF25-FLAG (Fig. 5a) and human HERC5 in transfected HEK293T cells (Fig. 5b) as analyzed by co-immunoprecipitation assays. To determine whether HERC5 is important for ORF25 ISGylation, we depleted HERC5 in HEK293T cells (Extended Data Fig. 5a) and analyzed ORF25 ISGylation. Precipitation under denaturing condition followed by immunoblotting showed that HERC5 depletion by two pairs of shRNA either abolished or greatly reduced ORF25 ISGylation (Fig. 5c). Under the same conditions, depletion of mouse HERC6 also abolished ORF25 ISGylation in MEFs infected with recombinant MHV68.ORF25-FLAG (Fig. 5d and Extended Data Fig. 5b). The MHV68.FLAG-ORF25-K1333R served as a negative control for this experiment. To further identify viral proteins that modulate ORF25 ISGylation, we performed affinity purification of ORF25 under physiological condition using MEFs infected with MHV68.FLAG-ORF25 and tandem mass spectrometry analysis to categorize the ORF25- binding proteins (Fig. 5e). In the ORF25-binding protein, ORF75c was identified as a major ORF25 interactor by tandem mass spectrometry (Fig. 5f). ORF75c has intrinsic E3 ligase activity to promote ubiquitination. We then assessed whether ORF75c can promote ORF25 ISGylation. In transfected HEK293T cells, ORF75c expression can enhance ORF25 ISGylation (Fig. 5g). This result suggests that ORF75c either acts as an E3 ligase or stimulates HERC5’s E3 ligase activity to promote ORF25 ISGylation. In HERC5- depleted HEK293T cells, ORF75c expression failed to enhance ORF25 ISGylation (Fig. 5h), supporting that ORF75c stimulates HERC5 to promote ORF25 ISGylation. Consistent with this, ORF75c interacted with HERC5 in transfected HEK293T cells by co-immunoprecipitation assay (Fig. 5i) and its expression increased ORF25 interaction with HERC6 (Fig. 5j). Taken together, these results show that human HERC5 and mouse HERC6 are E3 ligases necessary for ORF25 ISGylation.

**Figure 5.**
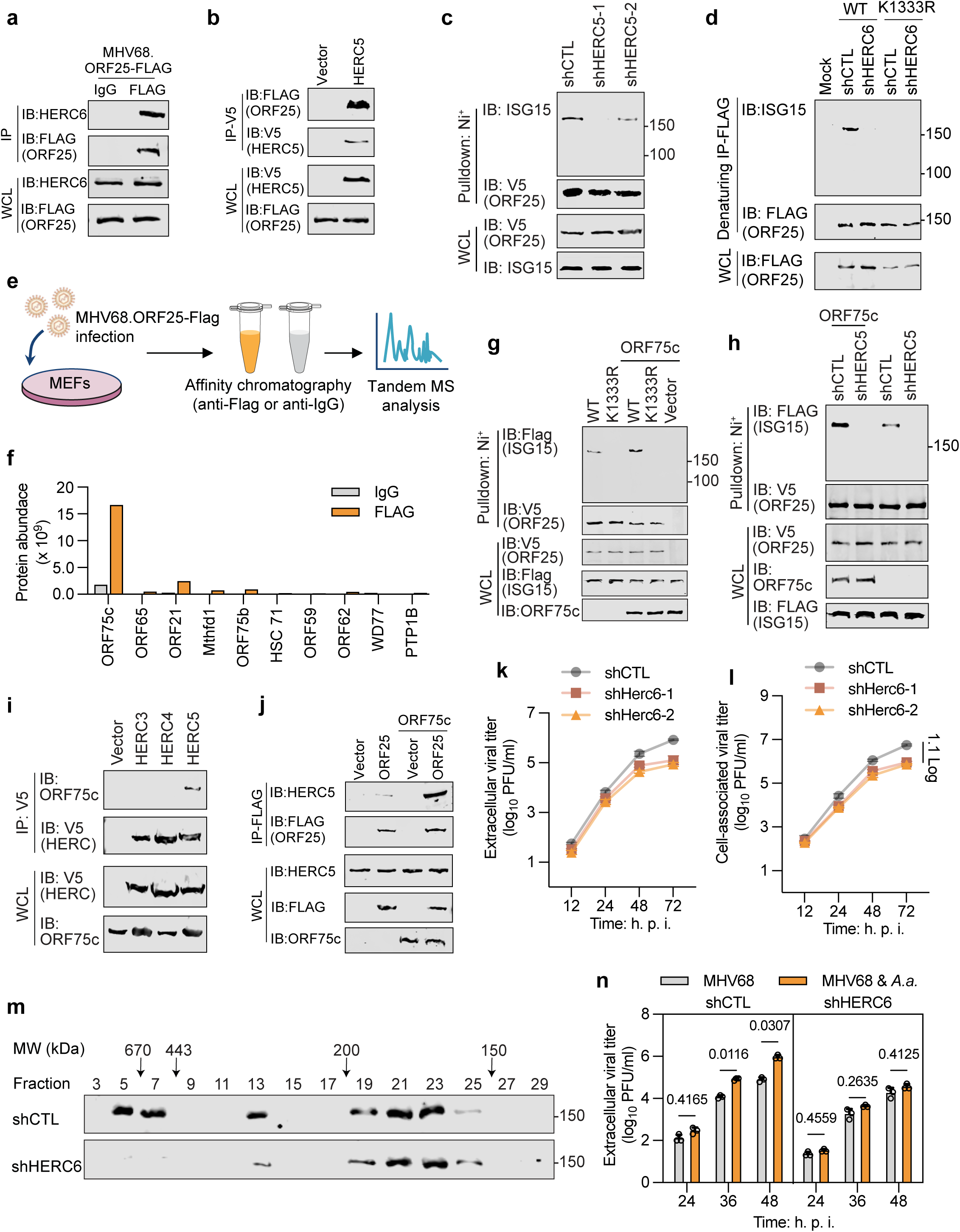
Mouse HERC6 and human HERC5 serve as an E3 ligase of the MHV68 ORF25. **a**,**b**, Immunoblotting of precipitated proteins and whole cell lysates (WCLs) from MEFs infected with MHV68.ORF25-FLAG (MOI=2, 48 hpi) (**a**) or HEK293T cells co-transfected with ORF25-FLAG and empty vector or human HERC5-V5 (**b**). **c**, shCTL and shHERC5 HEK293T cells were co-transfected with FLAG-ISG15 and ORF25-His/V5. ORF25 was purified with Ni-NTA agarose followed by immunoblotting with indicated antibodies. **d**, shCTL and shHERC6 MEFs were infected with MHV68.ORF25-WT-FLAG or MHV68.ORF25-K1333R-FLAG. ORF25 was immunoprecipitated with anti-FLAG M2 agarose under denaturing condition. Precipitated proteins and WCLs were analyzed by immunoblotting. **e**,**f**, Schematic illustration of the proteomics analysis of ORF25-binding proteins. MEFs were infected with MHV68.ORF25-FLAG (MOI = 2, 48 hpi). ORF25 was purified with anti-FLAG M2 agarose. Anti-IgG served as a negative control. Purified protein was trypsin digested and subjected to LC-MS/MS (**e**). ORF25-binding proteins identified by the proteomics analysis and the protein abundance were shown in the bar graph (**f**). **g**, HEK293T cells were co-transfected with FLAG-ISG15, ORF25-His/V5 (WT or K1333R), with or without ORF75c as indicated. ORF25 was precipitated with Ni-NTA agarose followed by immunoblotting with indicated antibodies. **h,** shCTL and shHERC5 HEK293T were co-transfected with FLAG-ISG15, ORF25-His/V5-WT, with or without ORF75c co-expression. ORF25 was precipitated with Ni-NTA agarose followed by immunoblotting with indicated antibodies. **i**, HEK293T cells were co-transfected with FLAG-ORF75c and HERC-V5. Co-immunoprecipitated protein and WCLs were analyzed by immunoblotting with indicated antibodies. **j**, HEK293T cells were co-transfected with ORF25-FLAG with or without ORF75c. Precipitated proteins and WCLs were analyzed by immunoblotting with indicated antibodies. **k**,**l**, Multi-step growth curve of MHV68 (MOI=0.01) in shCTL and shHERC6 MEFs was characterized by plaque assay using the extracellular virions (**k)** and infected cell lysates (**l**). **m**, ORF25 was purified in shCTL and shHERC6 MEFs infected with MHV68.ORF25-FLAG (MOI=2, 48 hpi) with anti-FLAG M2 agarose. Purified ORF25 was subjected to size exclusion chromatography and analyzed by immunoblotting with anti-FLAG. **n**, shCTL and shHERC6 MEFs were infected with MHV68 (MOI=0.1) for 2 h, then coinfected with or without *A.a.* as described in Fig. 1a. MHV68 titer in the medium at indicated time points was determined by plaque assay.

To assess the effect of HERC6 knockdown on MHV68 lytic replication, we first examined the multi-step growth curve of MHV68. We found that HERC6 knockdown reduced MHV68 yield by ∼10-fold at 48 and 72 hpi when extracellular infectious virions were measured by plaque assays (Fig. 5k). Similar effect was observed on intracellular virions, albeit to a less extent (Fig. 5l). Next, we examined ORF25 oligomerization in MEFs infected with MHV68 when HERC6 was depleted with shRNA. Consistent with the reduced viral replication, depletion of HERC6 also abolished the pentamer/hexamer species and greatly reduced the dimer species of ORF25 by size exclusion chromatography (Fig. 5m). Finally, we tested whether *A.a.* can stimulate MHV68 lytic replication when HERC6 was depleted. Plaque assay assessing extracellular infectious virions showed that HERC6 depletion diminished the replication of MHV68 that was elevated by *A.a.* co-infection (Fig. 5n). Collectively, these results show that HERC6 depletion recapitulates the phenotype of MHV68 replication in ISG15-deficient MEFs and that of the recombinant MHV68.ORF25-K1333R virus.

### ISGylation of herpesvirus capsid proteins is conserved

Herpesviruses share highly conserved capsid structure and plausibly mechanisms governing capsid assembly. Thus, we tested whether ISGylation-mediated capsid assembly is operating in the infection of other herpesviruses. We chose herpes simplex virus 1 (HSV-1), murine cytomegalovirus (MCMV), and Kaposi’s sarcoma-associated herpesvirus (KSHV) that represent α, β, and γ herpesviruses. To determine whether the major capsid protein UL19 of HSV-1 is ISGylated, we performed immunoprecipitation under denaturing condition using HeLa cells infected with recombinant HSV-1 carrying UL19-FLAG. Immunoblotting with anti-ISG15 antibody detected multiple species of UL19 (Fig. 6a). To probe the role of ISG15 in HSV-1 lytic replication, multi-step growth curve of HSV-1 infection in wildtype and *Isg15^-/-^* MEFs were performed, which showed that loss of ISG15 expression lowered extracellular viral titer by 0.5 to 1.5 orders of magnitude (Fig. 6b). Next, we purified UL19 from wildtype and *Isg15^-/-^* MEFs infected with HSV-1 carrying FLAG-UL19 and performed mass spectrometry to identify the sites carrying the GG motif. Comparison between sites modified by diglycine, we found that loss of ISG15 greatly reduced the frequency of modification by diglycine at K1041 and K1275 (Fig. 6c). Specifically, the frequency of diglycine modification at K1041 was reduced from ∼30% to zero and that of K1275 from ∼12% to 4% by ISG15 deficiency. Furthermore, the frequency of diglycine modification of K1041 and K1275 ranked the highest two. Interestingly, the frequency of diglycine modification at K224 and K373 was significantly elevated by ISG15 deficiency, which calls for further investigation of ISGylation-related post-translational modification of UL19. To assess the contribution of these two lysine residues to UL19 ISGylation, we generated recombinant HSV-1 carrying K1041R and K1275R mutations, designated HSV-1 2KR. Compared with wildtype HSV-1, HSV-1 2KR failed to give rise any ISG15 signal when analyzed by immunoprecipitation under denaturing condition and immunoblotting (Fig. 6d). Furthermore, the recombinant HSV-1 2KR replicated at 0.5 to 1 order of magnitude lower than wildtype HSV-1 (Fig. 6e). When extracellular released virions were analyzed by sucrose gradient centrifugation, we found that the HSV-1 2KR mutant generated more A- and B-type virions, and correspondingly less C-type virions (Fig. 6f and 6g). These results collectively support that HSV-1 UL19 is ISGylated and it is important for the virion maturation into C-type virions.

**Figure 6.**
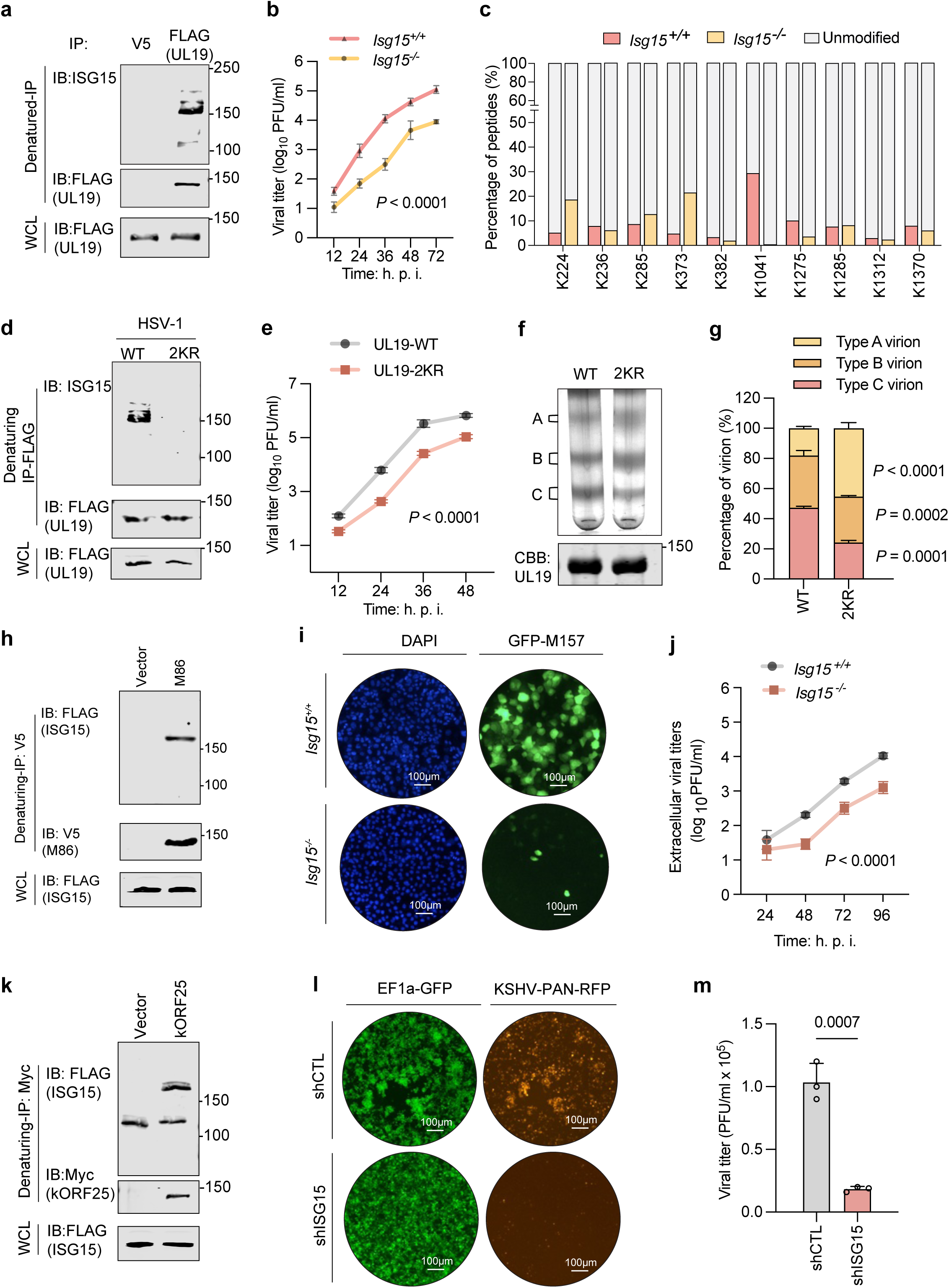
ISGylation of herpesvirus capsid proteins is required for efficient viral replication. **a**, Recombinant HSV-1 (HSV-1.UL19-FLAG) carrying the FLAG epitope at the C-terminus of UL19 was engineered using BAC. HeLa cells were infected with HSV-1.UL19-FLAG (MOI=2, 48 hpi). Precipitated proteins and whole cell lysate (WCL) were analyzed by SDS-PAGE and immunoblotting with indicated antibodies. **b**, Multi-step growth curve of HSV-1 (MOI = 0.01) in *Isg15^+/+^*and *Isg15^-/-^*MEFs was characterized by the plaque assay using extracellular virions in the medium. **c**, UL19 was purified from *Isg15^+/+^*and *Isg15^-/-^*MEFs infected with HSV-1.UL19-FLAG (MOI=2, 48 hpi) and analyzed by LC-MS/MS. ISGylation efficiency is the ratio of the abundance of GG-modified peptide to that of the total peptide for a specific site. Recombinant HSV-1 (HSV-1.UL19-2KR) containing K1041R and K1275R mutations was generated using BAC. **d**, HeLa cells were infected with HSV-1.UL19-WT and HSV-1.UL19-2KR (MOI=2, 48 hpi). UL19 was purified with anti-FLAG M2 agarose under denaturing condition and subjected to immunoblotting with indicated antibodies. **e**, Multi-step growth curve of indicated recombinant HSV-1 (MOI=0.01) in MEFs was characterized by plaque assay using extracellular virions. **f**,**g**, Virion assembly analysis of HSV-1.UL19-WT and HSV-1.UL19-2KR. HeLa cells were infected with indicated recombinant HSV-1 (MOI=0.1, 48 hpi). Extracellular virions in the medium were collected, concentrated by ultracentrifugation, and resolved by sucrose gradient centrifugation. The virion bands were photographed (**f**) and quantified by ImageJ (**g**). **h**, ISGylation of V5-M86 in transiently transfected HEK293T cells that expressed FLAG-ISG15, assessed by immunoprecipitation under denaturing condition with anti-V5 agarose followed by immunoblotting with indicated antibodies. **i**, *Isg15^+/+^* and *Isg15^-/-^* MEFs were infected with MCMV GFP-M157 (MOI=0.5) and examined by fluorescence microscopy at 72 h after MHV68 infection. Representative images were shown. Scale bars, 100 μm. **j**, Multi-step growth curve of MCMV (MOI=0.01) in *Isg15^+/+^*and *Isg15^-/-^*MEFs was characterized by plaque assay using extracellular virions. **k**, ISGylation of kORF25-Myc in transiently transfected HEK293T cells that expressed FLAG-ISG15, E1, and E2, assessed by immunoprecipitation under denaturing condition with anti-Myc antibody followed by immunoblotting with indicated antibodies. **l** and **m**, shCTL and shISG15 HCT116 cells were infected with KSHV (MOI=5, 48 hpi) and examined by fluorescence microscopy. Representative images were shown (**l**). Scale bars, 100 μm. Virus-containing medium was analyzed by plaque assay using HEK293T cells (**m**).

To determine whether MCMV major capsid protein M86 is ISGylated, we performed precipitation under denaturing condition using transfected HEK293T cells. As shown by immunoblotting, the ISG15 signal was readily detected in precipitated M86 (Fig. 6h). Furthermore, loss of ISG15 in MEFs reduced MCMV lytic replication as indicated by the fluorescence of the GFP-tagged M157 expressed from a recombinant MCMV and viral titer of the extracellular virions (Fig. 6i and 6j). Under similar conditions, we found that the KSHV ORF25 major capsid protein was ISGylated in transfected HEK293T cells (Fig. 6k). Depletion of ISG15 in HCT116 greatly reduced the productive replication of KSHV as indicated by RFP expression and extracellular infectious virions (Fig. 6l, 6m, and Extended Data Fig. 5c). These results concluded that the ISGylation of major capsid of MCMV and KSHV promotes their productive infection.

## Discussion

ISG15 is an IFN-stimulated gene whose protein product can be conjugated to other proteins, which modulates innate immune response^27, 28^. Emerging studies also demonstrate that ISG15 is secreted to extracellular compartment and acts on target cells to mediate immune response via binding to plasma membrane receptors^29, 30^. Both antiviral and proviral roles of ISG15 have been reported, though antiviral activities of ISG15 and ISGylation have been the focal point of research^31, 32, 12^. Though emerging, the proviral roles of ISG15 and ISGylation are less understood, and seemingly counterintuitive. Here, we report that MHV68 potently induces the expression of ISG15 and protein ISGylation to promote viral lytic replication. Mechanistically, ISG15 conjugation to the major capsid protein (ORF25) enables the oligomerization of ORF25 and capsid assembly of MHV68. The ISGylation-resistant K1333R mutation abolished ORF25 oligomerization and reduced virion assembly/maturation. The phenotype is reflected by the accumulation of A- and B-type virions, and a deficit in C-type virions, all of which were recapitulated by wildtype MHV68 infection in ISG15-deficient MEFs. The striking phenotypes of MHV68 lytic replication imposed by the K1333R and ISG15- deficiency are consistent with the exceedingly high frequency of GG-modified peptides containing K1333 in wildtype MEFs compared to that of *Isg15^-/-^*MEFs, with MHV68 infection. Excitingly enough, the neighboring H1332K mutation in the K1333R background restored most, if not all, of the ISGylation and oligomerization of ORF25, as well as virion maturation in producing C-type virions. These results unambiguously demonstrate the key role of ISGylation of ORF25 in MHV68 lytic replication via genetic, biochemical, and virological assays. Profiling the major capsid proteins of HSV-1, KSHV, and MCMV, we found that these proteins are ISGylated and ISGylation deficiency, either via depleting or deleting ISG15 expression or genetically engineered recombinant virus (HSV-1), reduced their lytic replication, suggesting that ISGylation constitutes a regulatory mechanism shared by multiple herpesviruses. In addition to ORF25, other virion structural proteins, such as M9 small capsid protein (ORF65), tegument (ORF75c) and portal (ORF43) proteins, were identified as ISGylation targets. The regulatory roles of ISGylation in physiological function of these proteins await further investigation. Interestingly, a previous study reported the antiviral activity of ISG15 in human cytomegalovirus (HCMV) infection^33^. The reasons underlying these seemingly paradoxical findings are not clear, although we did not examine the effect of ISG15 on HCMV infection due to the slow kinetics of productive infection. Additionally, loss of ISG15 rendered mice more susceptible to MHV68 infection, supporting the *in vivo* antiviral activity of ISG15^34^. Although ISG15 is classically viewed as an antiviral effector, accumulating evidence indicates that its role is highly virus-specific and that ISG15 or ISGylation can be co-opted by certain viruses, including HCV, HBV, and vaccinia virus, to promote viral replication or release^35, 36, 37^. Our study has identified a proviral activity of ISG15 intrinsically within MHV68-infected cells, which was suggested by a recent study. The antiviral activity of ISG15 in mice may rest on the immune response that requires ISG15. As such, the antiviral activity of ISG15 against MHV68 is likely underestimated in a global deletion mouse model. Further investigation is necessary to tease out the proviral and antiviral activity of ISG15 in MHV68 infection using mice.

Late steps of herpesvirus lytic replication, such as capsid assembly and tegument protein incorporation, are poorly characterized due to the limited availability of tools and reagents^7, 38^. The identification of ISG15 and viral protein ISGylation that are important for late stages of lytic replication of MHV68 and other herpesviruses, offers a platform that enables the dissection of regulatory events in capsid assembly and virion maturation. When extracellular virions were examined by sucrose gradient centrifugation, we found that ISG15 deficiency and the ISGylation-resistant K1333R mutation induced accumulation of A- and B-type virions and correspondingly reduced the infectious C-type virions of MHV68. These results point to the role of ISGylation in facilitating the proper recruitment and subsequent removal of the chaperone protein (ORF17) of the major capsid proteins and enabling successful viral DNA packaging. Consistent with this model, ISGylation of ORF25 promotes its oligomerization, thereby supporting capsomer formation and capsid assembly. Coincidentally, the functional role of ISGylation in potentiating protein oligomerization has been appreciated in other biological contexts, further supporting this mechanism. Concomitant to this process, ISG15 is likely removed from ORF25 within the assembled virion capsid. Though the ISGylation of K1333 was detected at the frequency of ∼80% in MHV68-infected cells, it was under detection limit using purified MHV68 virions. These observations collectively support that the capsid assembly and subsequent DNA packaging, as well as ORF25 ISGylation, are dynamically regulated during MHV68 lytic replication. Beyond the sites of ISGylation, including K1333, K910 and K729, additional lysine residues carrying the GG remnant of trypsin digestion were abundantly detected throughout the full-length ORF25, suggesting potential regulation by ubiquitination and SUMOylation in capsid assembly and virion maturation. This study reveals opportunities to leverage post-translational modification events to interrogate late-stage regulatory mechanisms in herpesvirus lytic replication and cellular events alike.

## Materials and Methods

### Cell Culture

HEK293T, NIH 3T12, HeLa, Vero, and HCT 116 cells were purchased from ATCC. Wildtype and *Isg15^-/-^* mouse embryonic fibroblasts (MEFs) were kindly provided by Dr. Lilliana Radoshevich (University of Iowa, USA). All cells were cultured in DMEM (HyClone) supplemented with 10% heat-inactivated FBS (Gibco) and 1% penicillin-streptomycin (Corning) and maintained in a humidified incubator with 5% CO2 at 37 °C.

### Plasmids

Point mutations of ORF25 and ISG15, including ORF25-K1333R (HR), ORF25- H1332K&K1333R (KR), and ISG15-G155A were generated by site-directed mutagenesis and confirmed by sequencing. Lentiviral expression constructs containing FLAG-ISG15 or FLAG-ISG15-G155A were generated from pCDH-CMV-EF1-Puro by molecular cloning. A cDNA construct was used to amplify and clone M86 into mammalian expression vectors. For the shRNA-mediated knockdown of HERC6, two pairs of synthetic DNA oligos were annealed and cloned into pLKO.1 (Sigma) that was digested with *Age*I and *EcoR*I. The pLKO.1 expressing the scrambled shRNA was purchased from Sigma.

### Lentivirus-mediated Stable Cell Line Construction

Lentivirus production was carried out in HEK293T cells. Briefly, HEK293T cells were co-transfected with packaging plasmids (pMD2.g and psPAX) and pCDH lentiviral expression vector or lentiviral shRNA plasmids. At 48 hours post transfection (hpi), the medium was harvested and filtered through a 0.22 μm membrane for further experiments.

To generate the reconstituted cell lines, lentivirus carrying FLAG-ISG15 or FLAG-ISG15-G155A was added to *Isg15*^-/-^ MEFs for spinfection with 8 μg ml^−1^ polybrene at 1,000g for 45 min. At 48 h post infection, puromycin was used to select cells at a final concentration of 2 μg ml^−1^. The expression of ISG15 was validated by western blot with antibodies to ISG15 or the FLAG epitope. All lentivirus-mediated stable cell line constructions were following this protocol unless specified.

### Viruses

MHV68 K3/GFP was used as previously described^39^. MHV68 K3/GFP were amplified in NIH3T12 cells, and viral titer was determined by a plaque assay in a NIH 3T12 monolayer. Briefly, a ten-fold serial-diluted virus-containing medium (FBS-free DMEM) was added to NIH 3T12 cells. For the viral titer of cell lysates, three rounds of freezing and thawing was applied before loading the sample. After 2 h of incubation, DMEM containing 5% FBS and 1% methylcellulose (Sigma-Aldrich) was overlaid to infected NIH 3T12 cells after removing the supernatant for the plaque formation. At 5 days post infection, NIH 3T12 monolayers were stained with 1% crystal violet for plaque counting. For MHV68 infection, cells were seeded in 6-well or 12-well plates one day prior to infection. On the day of infection, viral inocula were adjusted according to the desired MOI and cell confluency. Cells were incubated with the virus for 2 hours to allow viral entry, after which the inoculum was removed and replaced with fresh medium. Samples were collected at the indicated time points for downstream analyses. MHV68 K3/GFP was used for the MHV68 infection unless specified otherwise.

HSV-1and KSHV production and titration were performed as previously described^40,22^. MCMV was generated as previously reported^41^.

### MHV68 and Aggregatibacter actinomycetemcomitans (A. a.) Co-infection

*A. a.* strains D7SS glycerol stock was plated on modified trypticase soy broth (TSBYE) containing 3% trypticase soy broth, 0.6% yeast extract, and 1.5% agar plates and cultured at 37 °C with 5% CO₂ for 48 h. Bacterial colonies were isolated from plates and resuspended in antibiotic-free DMEM to prepare bacterial suspensions. The optical density at 600 nm (OD_600_) was measured to determine bacterial concentration, and infections were performed at the indicated Colony-Forming Unit (CFU).

For the coinfection, MEFs were seeded in 24-well plates and cultured in DMEM supplemented with 10% FBS but without penicillin or streptomycin at 37 °C in 5% CO₂ for 24 h. Cells were first infected with MHV68 (MOI = 0.1) for 2 h in antibiotic- and serum-free DMEM under the same incubation conditions. After infection, the medium was removed and replaced with fresh DMEM containing 10% FBS (without antibiotics), followed by infection with *A.a.* D7SS at 75 CFU/cell. Infected cells were incubated for 12 h, then the medium was replaced with fresh DMEM containing 10% FBS (without antibiotics) and further incubated. At the indicated time points, GFP intensity from MHV68-K3/GFP was recorded using a fluorescence microscope. Both cell-associated and extracellular viral titers were determined by plaque assay. Cells were also collected for quantitative RT-PCR and western blot analysis.

### RNA Extraction, Reverse Transcription, qRT-PCR, and DNA Extraction

Total RNA was extracted using TRIzol Reagent (Invitrogen). Approximately 1 μg total RNA was used for reverse transcription with M-MLV Reverse Transcriptase (Promega) according to the manufacturer’s instruction. The synthesized cDNA was used as a template in quantitative real-time PCR (qRT-PCR) reactions with SYBR master mix (Applied biosystems). Primers for q-PCR can be found in Table S1. Additional primer sequences may be found from previous publications.

For DNA extraction, cells were resuspended in 500 μL of NK buffer (50 mM Tris, 50 mM EDTA, 1% SDS, pH8.8), followed by the addition of 2 μL of 20 mg/mL proteinase K. The mixture was incubated at 55 °C for 2 h to digest proteins, then thoroughly vortexed and centrifuged at 300 × g for 2 min. The supernatant was carefully collected, and 170 μL of 7.5 M ammonium acetate was added, followed by vigorous vertexing to mix. Samples were centrifuged at 12,000 rpm for 10 min, and the resulting supernatant was transferred to a new microcentrifuge tube. An equal volume of isopropanol was added, and the mixture was inverted several times to mix thoroughly before centrifugation at 15,000 rpm for 10 min. The supernatant was discarded, and the DNA pellet was washed once with 1 mL of 75% ethanol. The resulting DNA pellet was dissolved in DNase-free water, quantified with a Nanodrop unit, and stored for downstream applications. To determine the viral genome copy number, plasmids expressing ORF50 and ORF57 were serially diluted from 10^-1^ to 10^-10^ and subjected to real-time PCR to obtain Cq values. The corresponding DNA copy numbers were calculated using a DNA Copy Number and Dilution Calculator. A standard curve correlating Cq values with copy numbers was generated and used to convert the Cq values of samples by real-time PCR into absolute genome copy numbers. Primers are described in Supplementary Table 1.

### Viral Entry Assay

*Isg15*^+/+^ or *Isg15*^-/-^ MEFs were infected with MHV68 (MOI = 5) for up to 2 h. Infected cells were disassociated by trypsin digestion and washed extensively with PBS. Total genomic DNA was extracted, and the MHV68 genome was quantified by real-time PCR using primers specific to MHV68 genes, such as ORF50 and ORF57. To perform the entry assay for the MHV68 virions containing ISGylation-resistant mutation (K1333R), MHV68.ORF25-K1333R progeny virions were collected and normalized by quantitative PCR for the packaged viral genome. Equal amounts of the viruses were used for entry assay as described above.

For progeny generated from *Isg15^-/-^* MEFs, viral input was normalized based on packaged genome copy number determined by quantitative real-time PCR.

### Virion Trafficking Assay

*Isg15*^+/+^ or *Isg15*^-/-^ MEFs were infected with MHV68 (MOI = 5) for up to 2 h. Infected cells were washed extensively with PBS, resuspended in fractionation buffer (20 mM HEPES, 10 mM KCl, 2 mM MgCl2, 1 mM EDTA, 1 mM EGTA, 1 mM DTT, 0.2% saponin and protease inhibitor cocktail), and incubated on ice for 15 min. Then, the cell suspension was passed through a 27 Ga needle for 10 times using a 1 ml syringe and left on ice for 20 minutes. After that, cell lysates were centrifuged at 720 x g (3,000 rpm) for 5 min. The pellet and supernatant were collected separately. The pellet contains nuclear DNA, while the supernatant contains cytosolic DNA. Each fraction was split for real-time PCR for extracted DNA and immunoblotting analysis.

### Virion Assembly Analysis

*Isg15*^+/+^ and *Isg15*^-/-^ MEFs were infected with MHV68 (MOI = 0.1). At 72 h post infection, the virus-containing medium was collected and centrifuged at 4,000 rpm for 20 minutes at 4 °C to remove cell debris. The supernatant was then subjected to ultracentrifugation at 32,000 rpm for 2 h at 4 °C using a Beckman Type 45 rotor to pellet virions. The resulting virion pellets were resuspended in TNE buffer (0.01 M Tris-HCl, 0.5 M NaCl, 1 mM EDTA, pH 7.5). Resuspended virions were loaded onto a 20-50% (w/v) sucrose gradient and centrifuged at 80,000 × g for 2 h at 4 °C in a Beckman SW41 rotor^42^. A Coomassie Brilliant Blue staining of the major capsid protein was performed to verify equal loading of virion samples prior to gradient analysis. Virions were visualized as milky bands under scattered light. Images of the gradient were captured, and the integrated density of each virion band was quantified using ImageJ software. The relative proportion of each band was calculated as a percentage of the total integrated density.

### Virion Release Analysis

*Isg15*^+/+^ and *Isg15*^-/-^ MEFs were infected with MHV68 (MOI = 2) for indicated time points. Extracellular virions were collected from the medium, and intracellular virions were released by subjecting pelleted cells to three rounds of freeze-thaw. Plaque assay was conducted using NIH 3T12 cells.

### Proteomics Analysis for ISGylated Peptides and Proteins

To identify ISGylated proteins in MHV68 infection, FLAG-ISG15-MEFs were infected with MHV68 (MOI = 2) for 48 hours. Affinity purification with anti-FLAG or anti-IgG agarose was performed under denaturing condition (boiling in 1% SDS) to ensure that only covalently bound proteins were precipitated. After overnight immunoprecipitation, on-bead trypsin digestion was performed. The beads were washed three times with the RIPA buffer followed by washing with 100 mM ammonium bicarbonate (Ambic). Samples were denatured, reduced with dithiothreitol (DTT), alkylated with iodoacetamide (IAA), and digested with trypsin (G-Biosciences) at 37°C overnight with gentle shaking. Digested peptides were acidified and eluted with 1% Trifluoroacetic Acid. Digested peptides were further desalted using a C18 tip (P200 Toptip, PolyLC) following the manufacturer’s instructions and then dried in a rotary evaporator. Purified peptides were re-suspended in 0.1% formic acid for liquid chromatography-tandem mass spectrometry (LC-MS/MS) analysis. Briefly, peptides separated with the C18 Acclaim PepMap column (75 μm id × 15 cm, 2 μm particle sizes,100 Å pore sizes, Thermo Scientific) were ionized at 1.9 kV in the positive ion mode. MS1 survey scans were acquired at the resolution of 70,000 from 350 to 1800 m/z, with a maximum injection time of 100 ms and AGC target of 1e6. MS/MS fragmentation of the 14 most abundant ions were analyzed at a resolution of 17,500, AGC target 5e4, maximum injection time 65 ms, and normalized collision energy of 26. Dynamic exclusion was set to 20 s and ions with a charge of +1, +7 and >+7 was excluded. MS/MS fragmentation spectra were searched with Proteome Discoverer SEQUEST (version 2.4, Thermo Scientific) against the in silico tryptic digested Uniprot all-reviewed Homo sapiens database (release June 2017, 42,140 entries). The maximum missed cleavages were set to 3. Dynamic modifications were set to oxidation on methionine (M, +15.995 Da), GG-modification on lysine (K, +114.042927 Da). Carbamidomethylation on cysteine residues (C, +57.021 Da) was set as a fixed modification. The maximum parental mass error was set to 10 ppm, and the MS/MS mass tolerance was set to 0.03 Da. The false discovery threshold was set strictly to 0.01 using the Percolator Node validated by q-value. The relative abundance of parental peptides was calculated by integration of the area under the curve of the MS1 peaks using the Minora LFQ node. This protocol was applied for all proteomics analysis unless specified otherwise.

To identify ISGylation sites of ORF25 using ISG15-WT or ISG15-G155A, HEK293T cells were co-transfected with Flag-ISG15-WT or Flag-ISG15-G155A together with ORF25-His/V5 for 12 h, followed by infection with MHV68 (MOI = 2) for 48 h. ORF25 was purified using anti-V5 agarose under denaturing condition (boiling in 1% SDS) and subjected to on-bead trypsin digestion. Peptide processing and LC-MS/MS analysis were performed as described above. Glycine-alanine (GA) modified lysine residue (K, +128.11859 Da) was set as a variable modification to identify ISG15-G155A-derived ISGylation sites. Spectral annotation was generated by the Interactive Peptide Spectral Annotator (IPSA, http://www.interactivepeptidespectralannotator.com/).

To identify ISGylation sites of ORF25 by ISG15-deficient MEFs, *Isg15^+/+^* and *Isg15*^-/-^ MEFs were infected with MHV68.ORF25-FLAG (MOI = 2) for 48 h. ORF25 was purified by anti-FLAG M2 agarose under denaturing condition and analyzed by LC-MS/MS. ISGylation at individual lysine residues was quantified by calculating the ratio of di-glycine-modified peptide abundance to total peptide abundance at each site. Comparison between *Isg^+/+^* and *Isg15^-/-^* cells enabled the identification of *bona fide* ISGylation sites based on their selective loss in the absence of ISG15.

To identify ORF25-binding proteins, ORF25 and associated proteins were precipitated with anti-FLAG M2 agarose beads or anti-IgG from MEFs infected with MHV68.ORF25-FLAG (MOI = 2, 48 hpi). Reduction, alkylation, trypsin digestion, and proteomics analysis were performed as described above.

All raw proteomics data have been deposited in the Proteomics Identifications Database (PRIDE) under the accession number PXD071940.

### ISGylation Assay

HEK293T cells were transiently transfected with plasmids encoding Flag-ISG15, viral proteins (V5- and His-tagged), UBE1L, and UbcH8. 48 hours post transfection, 10% of cell suspension was used for direct immunoblotting analysis, and the remainder was collected and resuspended in the guanidine buffer (10 ml of 6 M guanidinium-HCl, 0.1 M Na_2_HPO_4_/NaH_2_PO_4_, 0.01 M Tris/HCl, pH 8.0, 5 mM imidazole and 10 mM β-mercaptoethanol). Approximately 20 ul HisPur™ Ni-NTA Resin (Thermo Scientific) were then added and lysates were rotated at room temperature (RT) for 4 h. The beads were successively washed for 5 min for each step at room temperature with the following buffers: 6 M guanidinium-HCl, 0.1 M Na_2_HPO_4_/NaH_2_PO_4_, 0.01 M Tris/HCl, pH 8.0 plus 10 mM β-mercaptoethanol; 8 M urea, 0.1 M Na_2_HPO_4_/NaH_2_PO_4_, 0.01 M Tris/HCl, pH 8.0, 10 mM β-mercaptoethanol; 8 M urea, 0.1 M Na_2_HPO_4_/NaH_2_PO_4_, 0.01 M Tris/HCl, pH 6.3, 10 mM β-mercaptoethanol (buffer A) plus 0.2% Triton X-100; buffer A and then buffer A plus 0.1% Triton X-100. After the last wash His-tagged products were eluted by incubating the beads in 75 ml of 200 mM imidazole, 0.15 M Tris/ HCl pH 6.7, 30% glycerol, 0.72 M β-mercaptoethanol, 5% SDS for 20 min at RT. Samples were processed for immunoblotting or stored at −80 °C for further purposes.

For purifying ORF25 under denaturing condition, HEK293T cells or MEFs were infected with MHV68.ORF25-FLAG (MOI = 2) for 48 h. Cells were harvested and lysed in NP-40 lysis buffer (50 mM Tris–HCl, pH 7.4; 150 mM NaCl; 1% NP-40; 5 mM EDTA) supplemented with 20 mM β-glycerophosphate, 1 mM sodium orthovanadate, 10 mM N-ethylmaleimide, and 1% SDS. Lysates were sonicated and boiled, then diluted to a final SDS concentration of 0.1% before centrifugation to remove insoluble material. Supernatants were pre-cleared with protein CL4B agarose for 1 h at 4 °C and incubated with anti-FLAG M2 agarose for 4 h at 4 °C. Beads were washed extensively with RIPA buffer, and bound proteins were eluted by boiling at 95 °C for 10 min. Eluted proteins were resolved by SDS-PAGE and analyzed by immunoblotting using primary antibodies at 1:1000 dilution and IRDye 800-conjugated secondary antibodies (Li-Cor) at 1:10,000 dilution. Signals were detected using the Odyssey infrared imaging system (Li-Cor).

### Co-immunoprecipitation (co-IP) and immunoblotting

Whole cell lysates were prepared with NP40 buffer (50 mM Tris-HCl, pH 7.4, 150 mM NaCl, 1% NP-40, 5 mM EDTA) supplemented with 20 mM β-glycerophosphate, 1 mM sodium orthovanadate, and a protease inhibitor cocktail (Roche). Whole cell lysates were sonicated and centrifuged. The supernatant was pre-cleared with protein A/G agarose for 1 h. Pre-cleared samples were incubated with indicated protein A/G agarose conjugated with antibodies for 4 h at 4°C. The agarose beads were washed extensively, and samples were eluted by boiling at 95°C for 10 min. Precipitated proteins were analyzed by SDS gel electrophoresis followed by immunoblotting.

All immunoblotting was performed using the following primary antibodies: ISG15 (Santa Cruz Biotechnology, #sc-166755, 1:200 dilution), β-Actin (Abcam, #ab8226, 1:5000 dilution), FLAG (Sigma, #F3165, RRID: AB_259529, 1:1000 dilution), V5 (Bethyl Laboratories, #A190-120A, RRID: AB_67586, 1:1000 dilution), Histone H3 (Santa Cruz Biotechnology, #sc-517576). HERC6 was provided by Dr. Kei-ichiro Arimoto (University of California, San Diego). Rabbit sera against vGAT (ORF75c) was described previously. Proteins were visualized by the Odyssey infrared imaging system (LI-COR) followed by incubation with IRDye800/680-conjugated secondary antibodies (1:10,000 dilution, LI-COR).

### Generating Recombinant Virus

Recombinant MHV68 and HSV-1 were generated using the Bacterial Artificial Chromosome (BAC)-based Lambda Red recombination system coupled with the *I-Sce*I homing endonuclease^25^. Lambda Red recombination utilizes the bacteriophage-encoded enzymes Exo, Beta, and Gam to mediate homologous recombination at the ends of linear double-stranded DNA. PCR products containing positive selection markers serve as suitable substrates for these enzymes, provided they bear 40-50 bp extensions homologous to the target sequence. The *I-Sce*I endonuclease recognizes an 18 bp sequence absent from the E. coli genome, allowing safe expression in bacterial cells. This recognition site is engineered on the pEP-KanS plasmid just outside of the positive selection marker. Upon cleavage by *I-Sce*I, intramolecular Red recombination between the duplicated flanking sequences removes the selection marker.

To generate recombinant viruses, MHV68 BAC DNA or HSV-1 strain 17 BACs were transformed into *E. coli* GS1783 cells, which express the Lambda Red recombinase system. PCR cassettes containing the desired mutation and the pEP-KanS-I-SceI selection cassette were generated using step-down PCR with primers containing homology to the pEP-KanS plasmid and 40 bp upstream and 20 bp downstream homology arms to the viral genome. PCR products were purified using the GeneJET PCR Purification Kit. The pEP-KanS-I-SceI cassette was inserted into BAC DNA by electroporation using 50 ng of purified PCR product. Following a 5-hour recovery period, the entire culture was plated on LB agar containing chloramphenicol and kanamycin and incubated overnight at 32 °C. Single colonies were inoculated into liquid LB medium containing chloramphenicol and kanamycin and grown overnight at 32 °C with shaking at 220 rpm. BAC DNA was isolated by isopropanol precipitation, and insertion of the pEP-KanS-*I-Sce*I cassette was confirmed by PCR using primers located 150 bp outside the mutation site, yielding an expected 1.4 kb product.

Removal of the pEP-KanS-I-SceI cassette was achieved by induction of I-SceI expression with 1% arabinose and activation of the Red recombinase at 42 °C. Subsequent PCR and Sanger sequencing verified successful marker excision and correct introduction of the mutation. BAC DNA was purified using the ZR BAC DNA Miniprep Kit. To rescue recombinant viruses, 2 μg of purified BAC DNA was transfected into HEK293T cells using Lipofectamine 3000. CPE was typically observed 4 days post-transfection. Cells were harvested and subjected to three rounds of freeze-thaw to produce the P0 virus stock. The P0 virus was then passaged twice in NIH3T12 or Vero cells to amplify the recombinant virus stock for subsequent experiments.

Viral titer of recombinant viruses was determined by quantifying packaged viral genome copy numbers using real-time PCR. A standard curve was generated using the wild-type virus, for which both plaque-forming units and packaged genome copy numbers were measured in parallel. Genome copy number of recombinant viruses was then converted to infectious titers based on this standard curve. Specifically, the titer of MHV68.ORF25-WT was determined by both plaque assay and quantitative PCR of packaged viral genomes. For MHV68.ORF25-K1333R, viral input was normalized with the packaged viral genome by real-time PCR.

### Size Exclusion Chromatography

Cells infected with MHV68.ORF25-WT, MHV68.ORF25-K1333R (HR), or MHV68.ORF25-KR were harvested in ice-cold NP-40 lysis buffer. ORF25 was purified using anti-FLAG M2 agarose for 4 h at 4 °C and eluted with 3×FLAG peptide. Eluted proteins were subjected to size-exclusion chromatography on a Superdex 200 Increase 10/300 GL column (Cytiva) equilibrated with PBS. Protein standards (Gel Filtration High Molecular Weight Calibration Kit, Cytiva) were run in parallel, including blue Dextran (2,000 kDa), thyroglobulin (669 kDa), apoferritin (443 kDa), β-amylase (200 kDa), alcohol dehydrogenase (150 kDa), and bovine serum albumin (66 kDa). Fractions (0.8 mL) were collected, and proteins were concentrated by trichloroacetic acid (TCA) precipitation. Concentrated samples were analyzed by SDS-PAGE followed by immunoblotting.

### Data Mining of RNA Sequencing

RNA-seq data was downloaded from the SRA database. Differential gene expression was analyzed using DESeq2 (v1.16.1), genes were annotated using KEGG and GO database. Original sequencing dataset is available at SRA database: SRR19792326, SRR19792325, SRR19792324, SRR19792321, SRR25823338^24^.

### Data Analysis

Statistical analysis was performed using GraphPad Prism software for unpaired two-tailed Student’s *t*-test or analysis of variance (ANOVA). *P*-values less than 0.05 were considered statistically significant.

## Acknowledgments

We thank Dr. Kei-ichiro Arimoto for providing the HERC6 antibody and Dr. Xinghong Dai for providing protocol to purify herpesvirus virion. Graphic design was created with BioRender.com, for which the authors possess a license. This study is partly supported by a generous startup fund from the Herman Ostrow School of Dentistry and grants from NIH (AI184716, CA285192, AI180537) to P.F.

