## supplemental figures for "Bacteria-herpesvirus interaction reveals a proviral role of ISGylation in promoting herpesvirus capsid assembly"

### Extended Data Figure 1

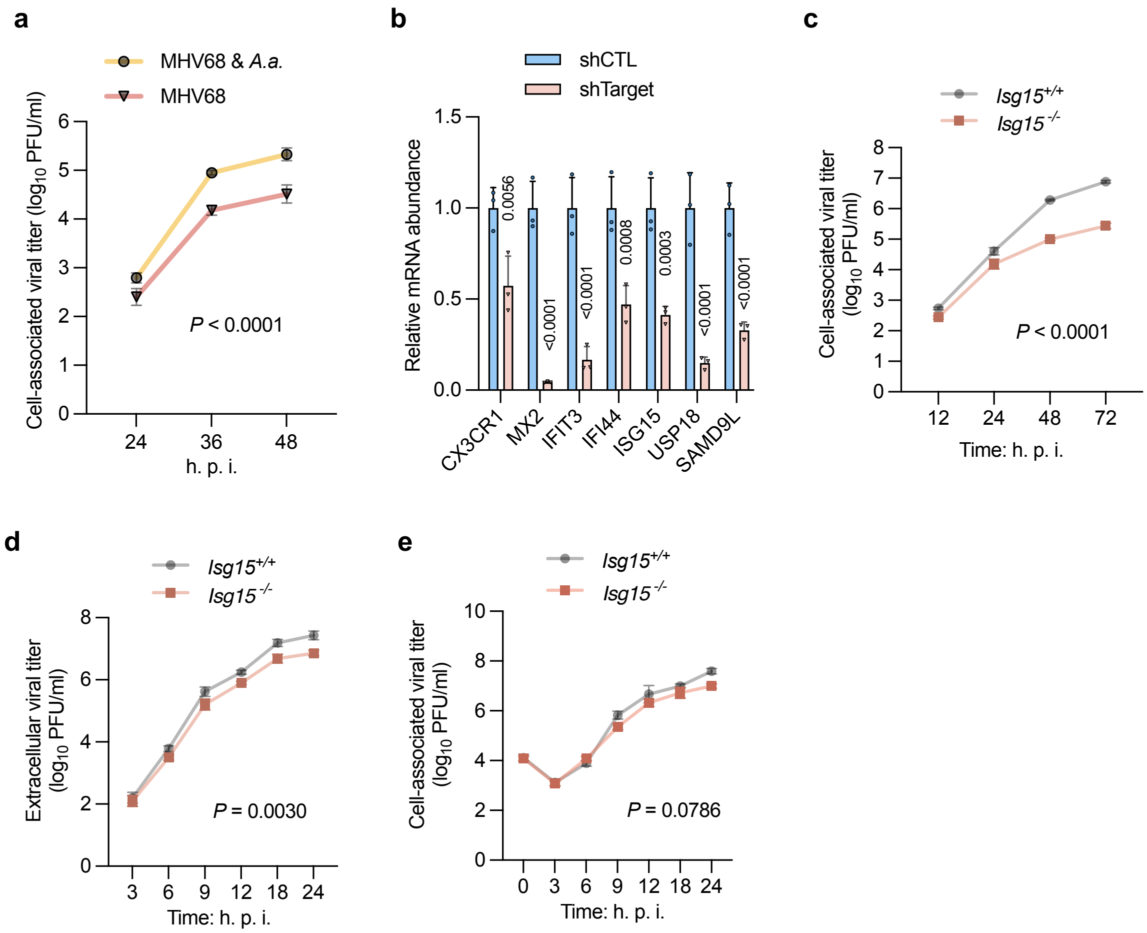

**Extended Data Figure 1**

**a,** MEFs were infected with MHV68 K3/GFP (MOI = 0.1. 2 hpi), with or without *A.a.* (75 CFU/cell, 12 hpi), washed with DMEM to remove *A.a.*, cultured with fresh medium. Cell-associated MHV68 titer at indicated time points was determined by plaque assay. **b**, Depletion of individual ISG was validated by reverse-transcription and real-time PCR using total RNA extracted from HEK293T cells infected with lentivirus-containing control (CTL) or ISG shRNA. **c**, Multi-step growth curve of MHV68 (MOI = 0.01) in *Isg15*$^{+/+}$ and *Isg15*$^{-/-}$ MEFs was characterized by plaque assay using infected cell lysates. **d,e**, Single-step growth curve of MHV68 (MOI = 5) in *Isg15*$^{+/+}$ and *Isg15*$^{-/-}$ MEFs was characterized by plaque assay using extracellular virions (**d**) and infected cell lysates (**e**).

#### Extended Data Figure 3

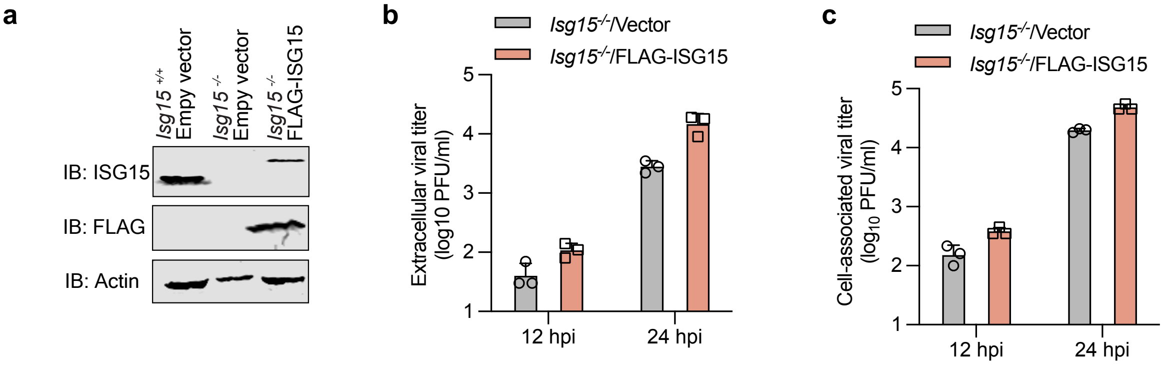


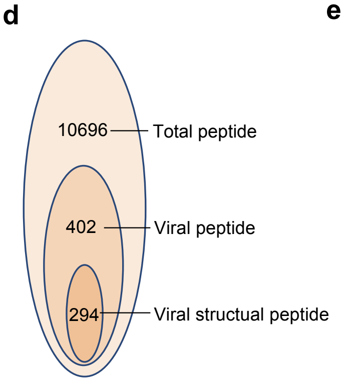


e

| Protein name | GG-Modified sites |
|---|---|
| Major capsid protein ORF25 | K24, K33, K94, K157, K259, K324, K373, K437, K573, K737, K910, K1067, K1215, K1333, K1344 |
| Tegument protein ORF75c | K422, K843 |
| Small capsid protein ORF65 | K7, K18, K34, K71/72, K135, K167 |
| Tegument protein ORF52 | K48, K78 |
| DNA replication processing factor ORF59 | K68, K158, K271, K305 |
| Capsid assembly protein ORF62 | K20, K193 |
| Triplex capsid protein 2 | K96, K293 |
| Tegument protein ORF75b | Not identified |
| Portal protein ORF43 | Not identified |

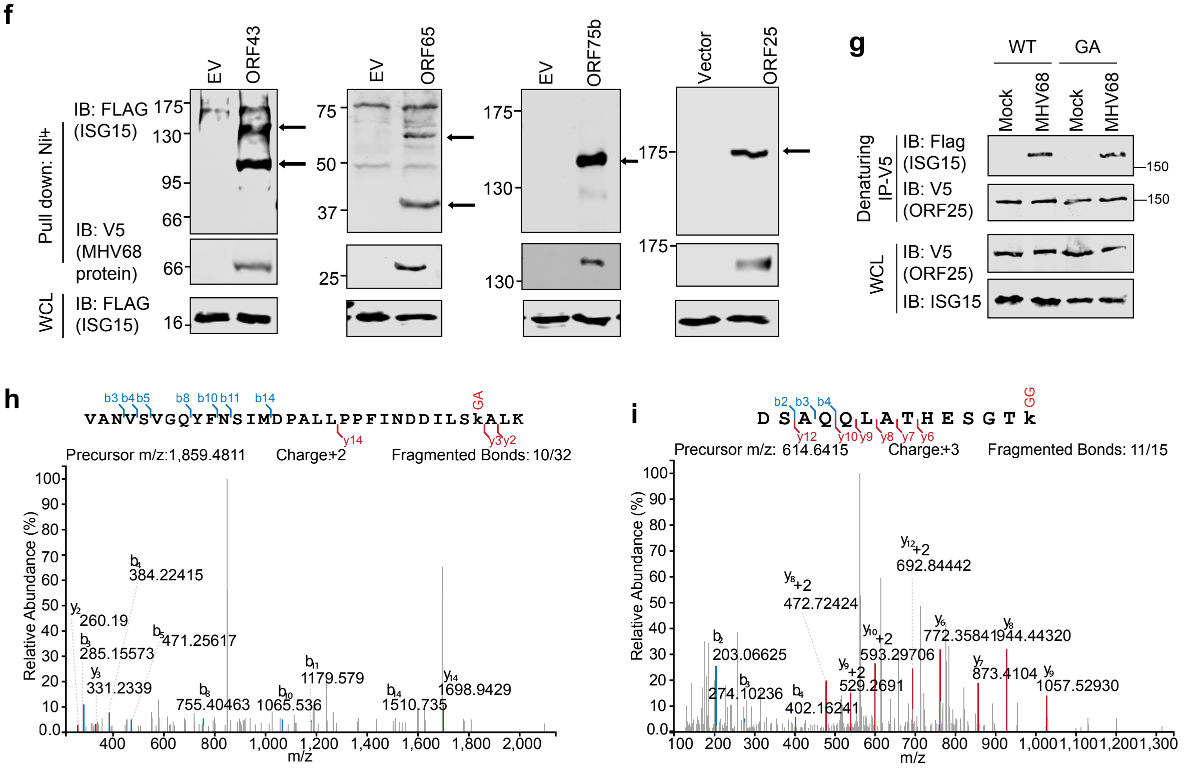

**Extended Data Figure 2**

**a**-**f**, Transcription of MHV68 genome templates in *Isg15*$^{+/+}$ and *Isg15*$^{-/-}$ MEFs at 4 hpi (**a**), 8 hpi (**b**), and 16 hpi (**c**) was analyzed by the reverse transcription and real-time PCR with gene-specific primers. The transcription of *ORF50* (**d**), *ORF59* (**e**), and *ORF75c* (**f**) at different time points in *Isg15*$^{+/+}$ and *Isg15*$^{-/-}$ MEFs was analyzed. **g**, Schematic illustration of the virion assembly analysis. Extracellular virions in the medium were collected and concentrated through ultracentrifugation at × 80,000 g for 2 h. The concentrated virions are resuspended in PBS and loaded on top of the sucrose gradient and centrifuged at 80,000 × g for 1 h. **h**, Viral release analysis was performed by measuring cell-associated and extracellular viral titer using plaque assay.

### Extended Data Figure 2

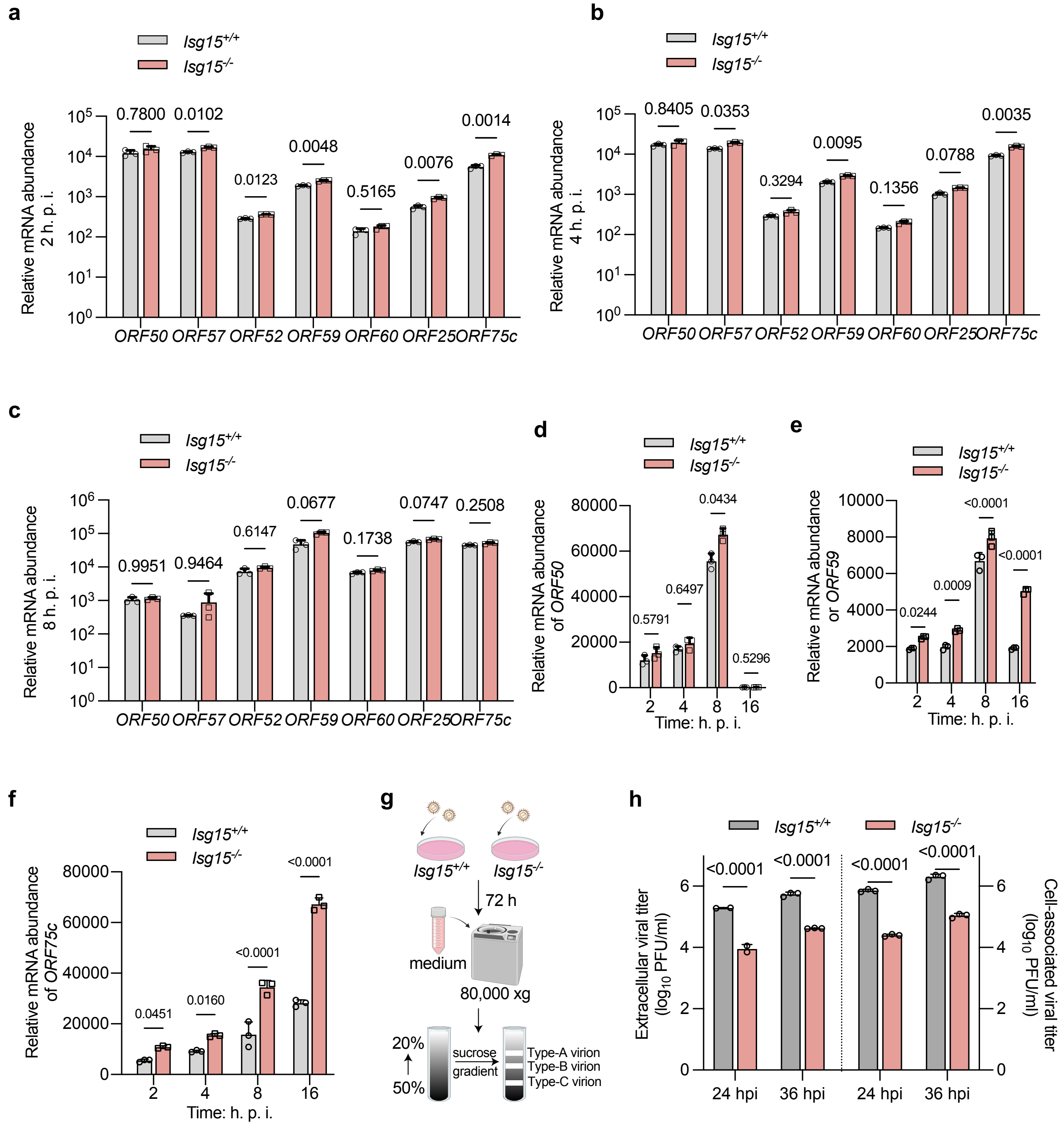

**Extended Data Figure 3**

**a**, *Isg15$^{-/-}$* MEFs were reconstituted with FLAG-ISG15 and were analyzed by SDS-PAGE and immunoblotting with indicated antibodies. **b**,**c**, Extracellular (**b**) and cell-associated (**c**) MHV68 titer in *Isg15$^{-/-}$*/empty vector and *Isg15$^{-/-}$*/Flag-ISG15 MEFs were determined by plaque assay. **d**, The pie chart showing the number of viral and cellular protein/peptides carrying GG motif. **e**, List of ISGylated proteins identified in the ISG15-conjugated-proteomic analysis. **f**, HEK293T cells were co-transfected with FLAG-ISG15, viral proteins-His/V5, E1, and E2. Precipitated proteins and whole cell lysates (WCLs) were analyzed by immunoblotting with indicated antibodies. **g**, HEK293T cells were transfected with FLAG-ISG15 (WT or G155A) and ORF25-His/V5, followed by MHV68 infection (MOI = 2, 48 hpi). ORF25 was precipitated with Ni-NTA agarose followed by immunoblotting with indicated antibodies. **h**, The m/z spectrum of the peptide containing K729 that carries the GA remnant. **i**, The m/z spectrum of the peptide containing K910 that carries the GG remnant.

#### Extended Data Figure 4

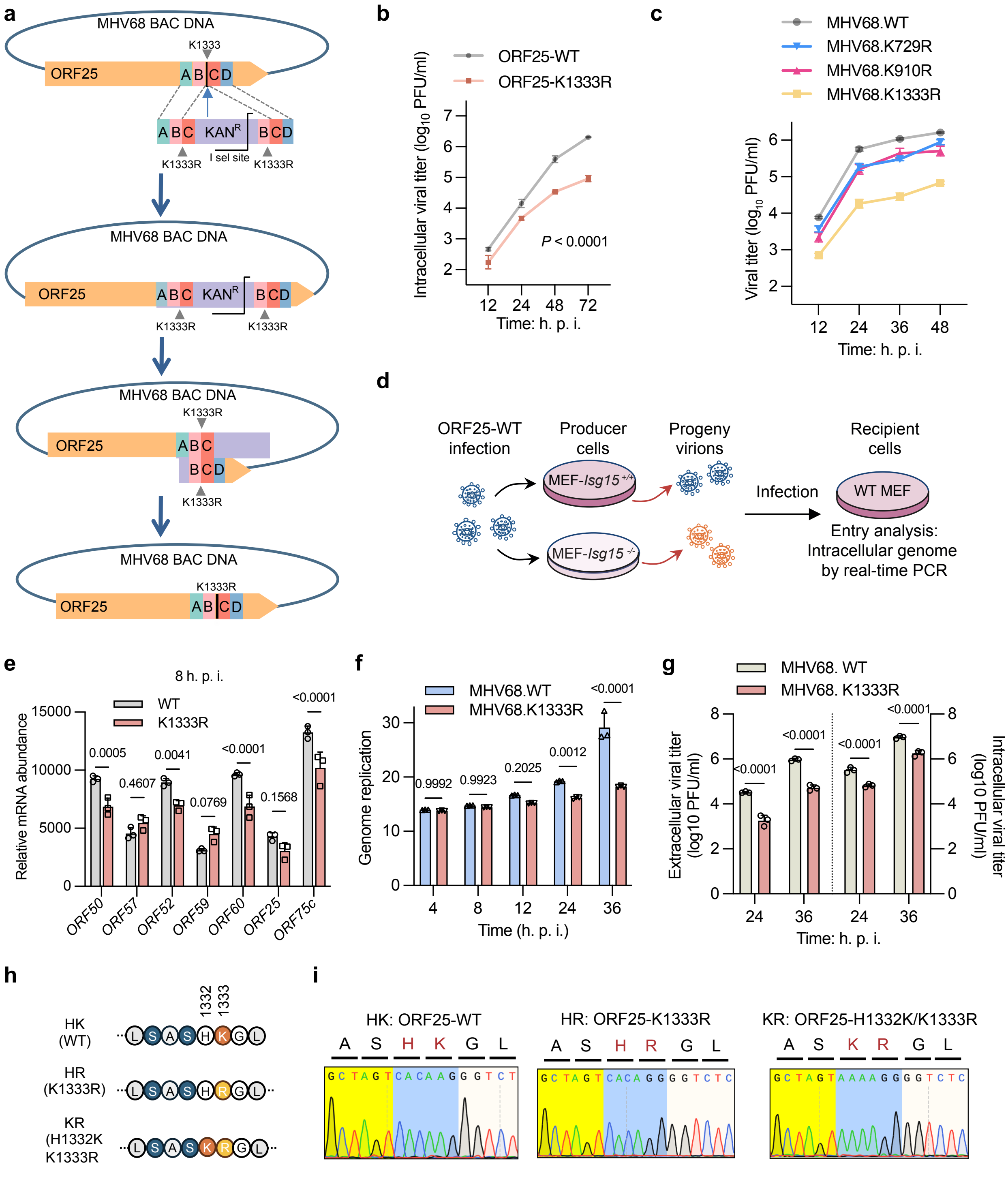

**Extended Data Figure 4**

**a**, Schematic illustration of generating the recombinant MHV68 carrying the ISGylation-resistant mutation. The diagram outlines each step of the recombineering workflow used to create the ORF25-K1333R mutation in MHV68 BAC. **b**, MEFs were infected with MHV68.ORF25-WT and MHV68.ORF25-K1333R. Viral input was normalized by packaged genome copy number. Multi-step growth curve of indicated recombinant MHV68 (MOI = 0.01) was characterized by plaque assay using infected cell lysates. **c**, MEFs were infected with MHV68.ORF25-WT, MHV68.ORF25-K1333R, MHV68.ORF25-K910R, and MHV68.ORF25-K729R. Multi-step growth curve of indicated recombinant MHV68 (MOI = 0.01) in MEFs was characterized by plaque assay using extracellular virions. **d**, Schematic illustration of entry analysis of the progeny virus. *Isg15*$^{+/+}$ and *Isg15*$^{-/-}$ MEFs were infected with MHV68 (MOI = 0.1, 72 hpi). Progeny virus was collected and normalized using the packaged genome copy number. Target cells was infected with the progeny virus (MOI = 5) and viral entry was determined by real-time PCR using total DNA. **e**, Transcription of MHV68.ORF25-WT and MHV68.ORF25-K1333R viral genome templates at 8 hpi was analyzed by the reverse transcription and real-time PCR with gene-specific primers. **f**, Genome DNA replication analysis. Total intracellular DNA was extracted from MHV68.ORF25-WT and MHV68.ORF25-K1333R infected MEFs and subjected to real-time PCR. **g**, Viral release analysis was performed by measuring intracellular and extracellular viral titers using plaque assays. **h**,**i**, Schematic illustration of recombinant MHV68 carrying the indicated mutations (**h**). Mutations engineered in the MHV68 genome were validated by sequencing (**i**).

### Extended Data Figure 5

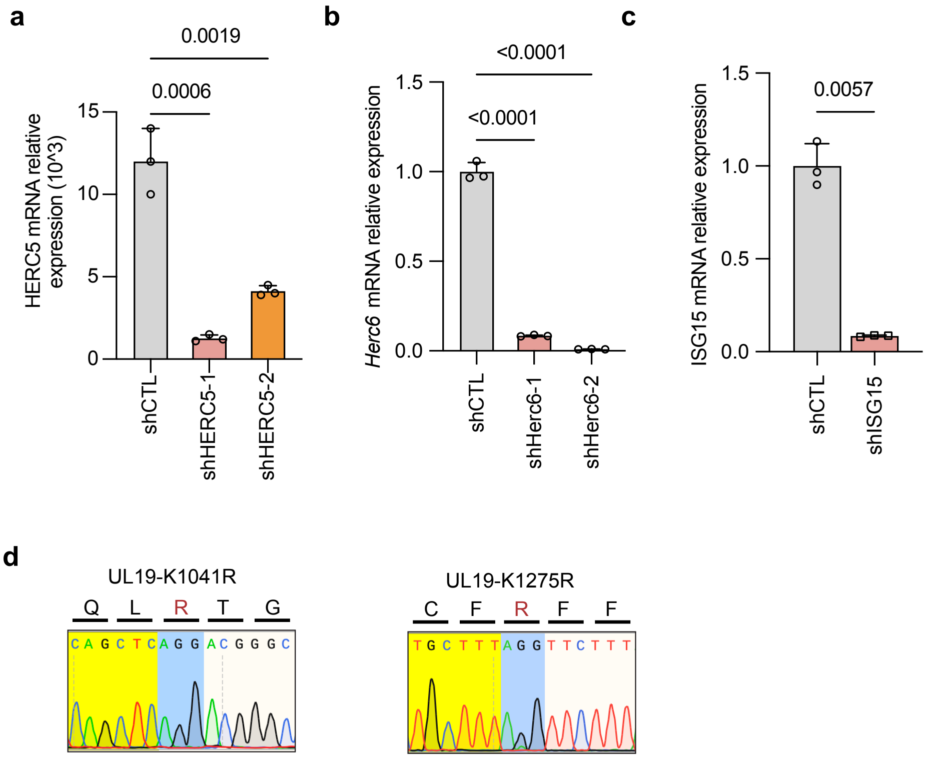

**Extended Data Figure 5**

**a**-**c**, HEK293T cells were transduced with lentivirus carrying shHERC5 (**a**), MEFs were transduced with lentivirus carrying shHERC6 (**b**), and HCT 116 cells were transduced with lentivirus carrying shISG15 (**c**). Knockdown efficiency was determined by reverse transcription and real-time PCR using total RNA. **d**, Mutations engineered in the HSV-1 genome were validated by sequencing.
